# Resting-state network topology characterizing callous-unemotional traits in adolescence

**DOI:** 10.1101/2021.08.27.457946

**Authors:** Drew E. Winters, Joseph T. Sakai, R. McKell Carter

## Abstract

**Background:** Callous-unemotional (CU) traits, a youth antisocial phenotype, are hypothesized to associate with aberrant connectivity (dis-integration) across the salience (SAL), default mode (DMN), and frontoparietal (FPN) networks. However, CU traits have a heterogeneous presentation and previous research has not modeled individual heterogeneity in resting-state connectivity amongst adolescents with CU traits. The present study models individual-specific network maps and examines topological features of individual and subgroup maps in relation to CU traits.

**Methods:** Participants aged 13-17 completed resting-state functional connectivity and the inventory of callous-unemotional traits as part of the Nathan Klein Rockland study. A sparse network approach (GIMME) was used to derive individual-level and subgroup maps of all participants. We then examined heterogeneous network features associated with CU traits.

**Results:** Higher rates of CU traits increased probability of inclusion in one subgroup, which had the highest mean level of CU traits. Analysis of network features reveals less density within the FPN and greater density between DMN-FPN associated with CU traits.

**Discussion:** Findings indicate heterogeneous person-specific connections and some subgroup connections amongst adolescents associate with CU traits. Higher CU traits associate with lower density in the FPN, which has been associated with attention and inhibition, and higher density between the DMN-FPN, which have been linked with cognitive control, social working memory, and empathy. Our findings suggest less efficiency in FPN function which, when considered mechanistically, could result in difficulty suppressing DMN when task positive networks are engaged. This is an area for further exploration but could explain cognitive and socio-affective impairments in CU traits.

## 1. Introduction

Callous unemotional (CU) traits describe an antisocial phenotype in youth (Frick & White, 2008) that represent the affective component of adult psychopathy (Barry et al., 2000; Frick et al., 2014). Adolescents with these traits incur a societal cost that is 10 times greater than typically developing youth (Cohen & Piquero, 2009; Foster et al., 2005). Because current treatments for antisocial phenotypes have limited efficacy (for meta-analysis: van der Stouwe et al., 2014), there is an ongoing need to develop a better understanding of these phenotypes. Identifying biomarkers related to callousness could inform development of new interventions and aid faster assessment of treatment response by monitoring treatment outcomes prior to changes being manifested behaviorally (Mayeux, 2004; Perez et al., 2014). However, this has been difficult because CU traits are heterogeneous (i.e., there are multiple variants underlying individual differences; e.g., Fanti et al., 2013; Fanti et al., 2018; Hadjicharalambous & Fanti, 2018; Catherine L. Sebastian et al., 2012). Prior investigations of neural mechanisms underlying CU traits use methods that do not account for individual heterogeneity. Given that adult psychopathy is viewed as a neurodevelopmental disorder with origins in youth (Frick & Viding, 2009) and children with CU traits share symptoms with adult psychopathy (Barry et al., 2000; Frick et al., 2014), it is critical to characterize neural networks, and previously uncaptured heterogeneity, underlying CU traits prior to adulthood. Thus, the present study examines CU traits amongst a community sample of adolescents using a novel method for characterizing neural networks that leverages individual heterogeneity.

Multiple lines of research document candidate brain abnormalities associated with CU traits and its adult counterpart, psychopathy (Barry et al., 2000). Task-based fMRI studies in adolescents suggest aberrant activity in the limbic, temporal, and frontal cortex regions in nodes associated with the salience (SAL), default mode (DMN), and frontoparietal (FPN; also called central executive) networks (Finger et al., 2008; Herpers et al., 2014; Jones et al., 2009; Lozier et al., 2014; Marsh et al., 2013; Pujol et al., 2012; Catherine L Sebastian et al., 2012; Veroude et al., 2016; Viding et al., 2012; White et al., 2012). Further support of the importance of these regions is found in studies measuring grey matter volume demonstrating abnormalities in similar regions (Caldwell et al., 2019; Cardinale et al., 2019; Cohn et al., 2016; Raschle et al., 2018; Rogers & De Brito, 2016; Sebastian et al., 2016; Wallace et al., 2014). These findings in youth are mirrored in adult studies on neural mechanisms underlying psychopathy (e.g., Sethi et al., 2018) supporting both the theory on psychopathy as a neurodevelopment disorder (Frick & Viding, 2009) as well as the importance of examining the SAL, DMN, and FPN.

As opposed to regional differences in volume or activation during a task, a recent theory by Hamilton et al. (2015) suggests that psychopathic traits involve an impaired integration within and between the SAL, DMN, and FPN. This dis-integration has been used to explain the prevalent theories on neural mechanisms underlying emotion (Blair, 2010; Kiehl, 2006) and attention (Larson et al., 2013; Neumann & Hare, 2008) impairments in adult psychopathy. Given the neurodevelopmental disorder view of psychopathy (Frick & Viding, 2009), dis-integration of these networks could also be applied to investigating impairments CU traits shares with psychopathy such as emotion (Blair, 2008), socio-affective functioning (Blair et al., 2005; Hawes & Dadds, 2012), and attention (Dadds et al., 2011). Initial support for this perspective includes both within and between network associations. For example, where we would expect stronger connectivity within network, CU traits associate with reduced connectivity within the SAL (Yoder et al., 2016) and DMN (Cohn et al., 2015; Umbach & Tottenham, 2020), as well as aberrant connectivity in the FPN (Cohn et al., 2015). Additionally, for between networks, where we would expect an anticorrelation between task positive and task negative networks in typically developing brains (Uddin et al., 2009), higher CU traits associate with a lower anticorrelation between the DMN and FPN (Pu et al., 2017). Although previous task-based findings provided insights into the brain regions activated by specific tasks, these results suggest etiology of CU trait impairments may reflect a trait like pattern of disintegration within and between these networks. This compelling body of work makes it clear that understanding the neural mechanism underlying CU traits could lead to novel intervention.

Several barriers remain before the neural mechanisms of CU can be used to drive new interventions. First, while these studies reveal similar differences in each network, the individual studies do not converge on regions or demonstrate the same network across all studies. This is possibly due to the heterogeneity of CU traits (e.g., Fanti et al., 2013; Fanti et al., 2018; Hadjicharalambous & Fanti, 2018; Catherine L. Sebastian et al., 2012) that is not modeled in the above adolescent studies. Ignoring the heterogeneity of CU traits can produce spurious connections that fail to describe the individuals in the sample, whereas modeling this heterogeneity can more accurately characterize network patterns in the brain (Gates & Molenaar, 2012). Given it has been demonstrated with psychopathic traits in adults (e.g., Baskin-Sommers et al., 2011; Dotterer et al., 2020; Efferson & Glenn, 2018; Espinoza et al., 2018; Korponay et al., 2017), it is critical we examine the heterogeneity of neural mechanisms underlying CU traits in adolescents.

Second, these studies often use methods that average across the entire times series to examine the strength of contemporaneous connections only. Ignoring lagged connections in resting state connectivity fails to model important neural connections – reflecting neuronal processes – critical for characterizing neural networks (Mitra et al., 2014). Modeling this as well as other important network function and architecture involves modeling the way networks are arranged, or their topology (De Vico Fallani et al., 2014). Network topology captures important features including relations between regions that have shown to be important for understanding health and mental health disorders (Rezaeinia et al., 2020; Stiso & Bassett, 2018) making it appropriate to leverage for understanding CU traits.

Third, most of this research is conducted on clinical or forensic samples when the spectrum of CU traits plausibly exists at some level (although lower) amongst community samples. The idea that psychiatric symptoms are on a spectrum and a disorder of neural circuits is consistent with the research domain criteria (RDoC) framework and a move away from symptom categories (Insel, 2014). Samples selected for extreme problem behaviors raise potential issues such as ceiling effects while also making it difficult to parse CU traits association with brain features from multiple other comorbid symptoms such as conduct problems. One exception is a study on a community sample children by Umbach and Tottenham (2020) that demonstrates the dimensionality of CU traits as well as its unique association with the brain independent of conduct problems. However, this study models the mean time series to examine the strength of contemporaneous connectivity, which may not identify important network features that are crucial for understanding brain associations with CU traits. Examining a community sample of adolescents using a topological approach that models individual variability can improve our understanding of CU traits by reducing uncertainty and endogeneity around associated brain correlates.

To address these methodological limitations, we use a novel network approach that includes traditional group level analyses while accounting for person-specific individual heterogeneity without averaging over the entire time series – group iterative multiple model estimation (GIMME; Beltz & Gates, 2017; Gates & Molenaar, 2012). GIMME uses data driven methods to, first, model statistically meaningful subgroup-level connections and, second, adds statistically meaningful connections at the individual-level that are unique to each participant (both are blind to CU traits). These connections include both contemporaneous and lagged connections. The results include the identification of subgroup patterns that model heterogeneity by providing person-specific estimates. GIMME outperforms other network modeling approaches, such as bayes nets and Granger causality, in simulation studies when modeling data heterogeneity (Gates & Molenaar, 2012). Studies examining psychopathology using GIMME have found variation in network patterns within the same diagnosis (Beltz, 2018; Price et al., 2017), including adult psychopathy (Dotterer et al., 2020), demonstrating reliable biological heterogeneity within diagnoses.

We can then examine topological features of person-specific networks in relation to CU traits, which capture more nuanced information beyond mean strength such as connection density, directionality, and centrality of node importance. Network density is the number of connections in a sparse network that indicate to what extent information travels between nodes within or between networks (De Vico Fallani et al., 2014). Node centrality is the number of connections into and/or out of a node which is used as a measure of node importance within a network or for communicating with other networks (De Vico Fallani et al., 2014; Kaiser, 2011). These network features have been shown to predict cognitive functioning by way of capturing information processing streams in the brain (Cohen & D’Esposito, 2016).

The present study aims to characterize network features of CU traits in a community sample of early-to-mid adolescents (ages 13-17) using person specific network connectivity within and between the SAL, DMN, and FPN. We used GIMME (Beltz & Gates, 2017; Gates & Molenaar, 2012) to generate person specific connectivity maps and to identify any subgroups of similar connectivity patterns. First, we examine CU traits association with identified subgroups. Then we examine associations of CU traits with individual-level network features of network density and node centrality across all participants. As suggested by previous research, we expect to see less connection density within the SAL, DMN, and FPN. For between network associations, given that adults demonstrate greater density (Dotterer et al., 2020) and adolescents demonstrate greater connectivity strength (Pu et al., 2017), we hypothesize greater density of between DMN-FPN connections in the present sample. We have no a priori hypotheses about subgroup homogeneity in network patterns, network centrality, or associations with CU trait subscales, but examine these to further characterize network features of these symptoms.

## 2. Methods

### 2.1. Sample

Adolescent participants (ages 13-17) were from the Nathan Kline Institute’s Rockland study, a study with a community sample on ages between 6-85 conducted in Rockland, New York (for study procedures see: Nooner et al., 2012). Data were collected from participants which involved a series of questionnaires and an fMRI session (both task and resting state). To our knowledge the resting state data and its relationship to CU traits in an adolescent sample have not been published.

### 2.2. Measures

#### Inventory of Callous-Unemotional Traits (ICU)

The ICU is a 24-item assessment of CU traits (Frick, 2004). The ICU has a confirmed factor structure and demonstrates convergent and divergent validity after removing two items for poor psychometrics (Kimonis et al., 2008). We used the same factor structure that excluded two items for poor psychometrics, which had adequate reliability in the present sample (α= 0.72). The ICU consists of three subscales: callousness, unemotional, and uncaring. Participants rate items on a four-point Likert scale from 0 (“not true at all”) to 3 (“definitely true”) on items such as “I do not show my emotions to others”. Higher scores indicate greater level of CU traits.

#### Covariates and Demographics

The youth self-report (YSR) is a measure for behavior problems in youth ages 11-18 (Achenbach & Rescorla, 2001). The externalizing (α= 0.87) subscale was used to control for conduct issues in the present analysis. Items from the externalizing subscale are rated on a three-point scale (0 not true – 2 very true) indicating how much they agree with the statement for the previous 6-months. Higher scores indicate higher externalizing symptoms. Validity and reliability of the YSR externalizing measure are within acceptable standards (Achenbach & Rescorla, 2001). Raw scores were used as recommended for research purposes by Achenbach and Rescorla (2001). We conducted analyses both controlled for and did not control for externalizing behavior to detect where CU traits accounted for unique variance.

Both pubertal development and sex were measured by the genital and breast development subscales of the Tanner assessment (α = 0.77), in which parents rated pictures representing development of secondary sex characteristics on a scale of 1 (pre-pubertal) to 5 (full maturity) (Petersen et al., 1988). Higher scores indicate greater developmental maturity. Given there is significant variation in the timing of puberty when measured by age (about five years, Parent et al., 2003) and that hormonal changes during puberty have a direct effect on the adolescent brain, which in turn impact mental state and behavior (Cameron, 2004; Dahl, 2004; Sisk & Foster, 2004), we choose to control for pubertal stage instead of using age. Similarly, sex effects are known to associate with CU traits and demonstrate differences in brain structure amongst adolescents with CU traits, we included sex as a covariate (Raschle et al., 2018).

#### Imaging Acquisition

Resting state images were analyzed from the Rockland dataset (www.nitrc.org/projects/fcon_1000/). We republish those parameters here for convenience. Images were collected with a Siemens TimTrio 3T scanner using a blood oxygen level dependent (BOLD) contrast with an interleaved multiband echo planar imaging (EPI) sequence. Participants were instructed to keep their eyes closed without falling asleep and to not think of anything while they let their mind wander. Each participant received an fMRI scan during resting state (260 EPI volumes; repetition time (TR) 1400ms; echo time (TE) 30ms; flip angle 65°; 64 slices, Field of view (FOV) = 224mm, voxel size 2mm isotropic, duration = 10 minutes) and a magnetization prepared rapid gradient echo (MPRAGE) anatomical image (TR= 1900ms, flip angle 9°, 176 slices, FOV= 250mm, voxel size= 1mm isotropic). T1 stabilization scan removal was not necessary given that the Siemens sequence collects images after saturation is achieved.

#### Resting-state fMRI preprocessing

Preprocessing was conducted using Statistical Parametric Mapping (SPM version 12; Penny et al., 2011) using the standard preprocessing pipeline via the CONN toolbox (version 18b; Whitfield-Gabrieli & Nieto-Castanon, 2012). Using the Artifact Detection Tools (ART; http://www.nitrc.org/projects/artifact_detect), motion outliers were flagged for correction if > 0.5mm and regressed out using binary motion covariates. No timing correction was used due to the short TR and multiband sequence used for acquisition. Physiologic CSF and white matter noise was regressed out of the BOLD signal using anatomic component-based noise correction method (aCompCor) (Whitfield-Gabrieli & Nieto-Castanon, 2012). Co-registered MPRAGE and EPI images were normalized to an MNI template. Smoothing of images were conducted using a 6mm Gaussian kernel. Finally, to preserve meaningful resting state associations, the data was bandpass filtered to between .008 and .09Hz (Satterthwaite et al., 2013).

#### Region of Interest Selection

Neural investigations on CU traits supported focus on the DMN, SAL, and FPN in the current analysis (Pu et al., 2017; Umbach & Tottenham, 2020; Yoder et al., 2016). Consistent with examining psychopathology generally (Menon, 2011), targeting these networks is in line with contemporary theory on psychopathic traits (Hamilton et al., 2015). Eight a priori core ROIs for each network were defined anatomically using the Harvard-Oxford atlas involving the medial prefrontal and posterior cingulate cortex (mPFC and PCC) for the DMN; anterior cingulate cortex (ACC) and bilateral insula for the SAL; as well as the bilateral lateral prefrontal cortices and bilateral posterior parietal cortices (LPFC and PPC) for the FPN (MNI coordinates: Supplementary Table 1). These ROIs represent the core regions of these networks and have been used to examine the associations within and between the DMN, SAL, and FPN (Menon, 2015; Menon & Uddin, 2010; Uddin et al., 2009).

#### GIMME

Network maps from each participants timeseries were constructed using R (Version 4.04; R Core Team, 2021) along with the ‘GIMME’ (Lane et al., 2021) and ‘lavaan’ packages (Rosseel, 2012). This method uses a data-driven sparse modeling approach that iteratively adds network connections and using LaGrange multipliers (Sörbom, 1989) to assess model fit and retain statistically meaningful connections (defined as connections that improve model fit for 75% of the sample). Sparse modeling used here minimizes spurious contemporaneous connections generated by saturated models (Gates et al., 2010). GIMME models contemporaneous (occurring at the same functional volume) or lagged (occurring at the previous volume) connections that apply to the entire sample, subgroups (data-derived subsample), or individual level (individual-specific connections). The result is a unified structural equation model (uSEM; Gates et al., 2011) for each participant that includes both contemporaneous and first order lagged connections. Non-significant connections are pruned during model fitting if their influence changed with the addition of new connections (Gates & Molenaar, 2012). Model-building ends when the network fits the data well, which were assessed using excellent fit criteria by Brown (2015) requiring two out of four alternative fit indices are met: root mean squared error of approximation (RMSEA)≤0.05, standardized root mean residual (SRMR)≤0.05, comparative fit index (CFI)≥0.95, or non-normed fit index (NNFI)≥0.95.

During model generation, GIMME identifies shared connectivity patterns while accounting for individual heterogeneity using a community detection algorithm (walktrap) (Gates & Molenaar, 2012), which simulations have proven to be a reliable method of detecting subgroups of network patterns (Gates et al., 2017; Pons & Latapy, 2005). Because each subgroup is defined by similarities in features, each is best described by their shared network features (e.g., Goetschius et al., 2020). Using both individual and subgroup level network features increase reliability of estimates for both individual and subgroups in comparison to other network approaches (Gates et al., 2017; Gates & Molenaar, 2012; Smith et al., 2011).

### 2.3. Statistical Analysis

All inferential statistical analyses were conducted with the statistical language R (Version 4.04; R Core Team, 2021). We extracted network features from the GIMME networks and conducted inferential statistic on these features using maximum likelihood estimation. Prior to path analyses, variables demonstrated linear relationships and data met assumptions for normality of residuals, auto correlation, and multicollinearity; and t tests revealed participants that were excluded were not significantly different from those included on demographics or variables of interest.

#### Network features

We extracted network features for all individual networks including density, node centrality within (SAL, DMN, and FPN) and between (DMN-SAL, DMN-FPN, and SAL-FPN) networks of interest. To account for individual differences, we extracted proportions and calculated positive and negative features separately. *Network density* involved the number of connections between nodes (regardless of contemporaneous or lagged connections) for within and between networks separately. *Node centrality* involved calculating the number of connections a node has relative to the number of potential connections within a network or between networks (connections / nodes – 1). Higher number of connections toward or away from a node or between nodes suggests greater information flow regarding that node or set of nodes.

#### Associations between network features and callous-unemotional traits

We first examined the probability of being included in identified subgroups given the presence of CU traits using a multinomial logistic regression. For individuals, we used path analysis to examine CU trait’s association with network features (network density and node centrality), which estimates all parameters simultaneously in one model and reduces multiple comparisons. Total CU traits were the independent variable of interest while controlling for sex and tanner stage. CU traits can be independent of conduct disorder (e.g., Baskin-Sommers, Waller, et al., 2015; Hyde et al., 2015), and, because the present analysis seeks to understand CU traits association with network features, we controlled for conduct problems. To ensure we are not capturing a suppression effect (e.g., Hyde et al., 2016; Lozier et al., 2014), we also ran all models without controlling for conduct problems. All results of were identical whether we controlled for conduct problems or not, suggesting there are no suppression or impact to model coefficients; thus, we only report on models that include conduct problems. To correct for multiple comparisons we used a false discovery rate correction (Benjamini & Hochberg, 1995; Benjamini & Yekutieli, 2001) for each analysis using ‘p.adjust’ in base R (R Core Team, 2021).

We then conducted exploratory and confirmatory analyses. First, to determine if any subgroups were more characteristic of CU traits, we used an ANOVA to examine if there were mean differences in CU traits across subgroups and conducted Tukey’s post hoc test for individual differences. Second, for individual-specific features, we ran exploratory analyses with CU trait subscales of callousness, uncaring, and unemotionality to determine if associations found with total scores were driven by these subscales. We did not control for multiple comparisons for exploratory or confirmatory analyses.

## 3. Results

### 3.1. Descriptives

To ensure integrity of the data, we excluded participants with a WAIS-II IQ score < 80 (α = .96; Wechsler, 2011), movement in global signal intensity or translation and rotational movement parameters > 3mm, or > 20% invalid scans. Out of a total of 122 participants between the ages of 13-17, 10 participants were removed for IQ < 80, 24 participants for motion, and four participants for invalid scans. Leaving a total of 84 participants for analysis. The analyzed sample were predominantly White (White= 63%, Black = 24%, Asian = 9%, Indian = 1%, other= 3%) balanced between sex (female = 45%) and a mean age of 14.59±1.48. Mean ICU total score (Boys= 23.11±7.56; Girls= 22.27±10.31; Total= 22.71±8.93) were within one standard deviation of other studies with community samples (Byrd et al., 2013; Essau et al., 2006).

### 3.2. GIMME

Resting state networks met a priori criteria of good fit on two out of four fit indices according to average fit (SRMR = 0.024, CFI = 0.953, see Supplementary Table 2) and identified significant heterogeneity indicated by low modularity (modularity = 0.024). Connections for all participants were detected between the lateral prefrontal cortex and posterior parietal cortex in the FPN and bilateral insulae in the SAL. Four subgroups were identified that comprised 65% participants (n=54). The remaining 35% of participants (n=30) did not match any subgroup (for subgroup depiction: Supplementary Figures 1 and 2). All person specific maps contained individual level connections (23.85±4) and had positive connections (16.49±2.04). Whereas 83.3% had negative connections (7.36±2.19). All models contained both contemporaneous and lagged connections (contemporaneous = 7.75±2.00, lagged = 16.18 ±2.17). Subgroup maps indicated connections were heterogeneous within three of the subgroups and homogeneous in the fourth subgroup only. Specifically, the first subgroup (n=19, modularity = 0.191) was characterized by 18 shared connections and 33 connections that varied between individuals; second subgroup (n=10, modularity = 0.190) was characterized by19 shared connections and 36 connections that varied between individuals; and the third subgroup (n=19, modularity = 0.225) was heterogeneous with 8 shared connections and 57 connections that varied between individuals, whereas the fourth subgroup (n=6, modularity = 0.213) was homogeneous with 24 shared connections and no individual connections (individual heterogeneity depicted in Supplementary Figure 3). The first subgroup had the highest overall average network density (Supplementary Table 3). A depiction of network features can be found in Supplementary Figure 2.

### 3.3. Subgroup Association with Callous-Unemotional Traits

Increases in CU traits associated with increases in odds of being in the heterogeneous second subgroup (*β=* 0.17, *p _(FDR corrected)_* = 0.048, odds ratio = 1.19; see Table 1 and Figure 1 Panel B.), which performed better than a null model (ΔChi= 25.14, p= 0.012). Associations were not significant with other subgroups and conduct problems had no significant associations.

**Figure 1.**
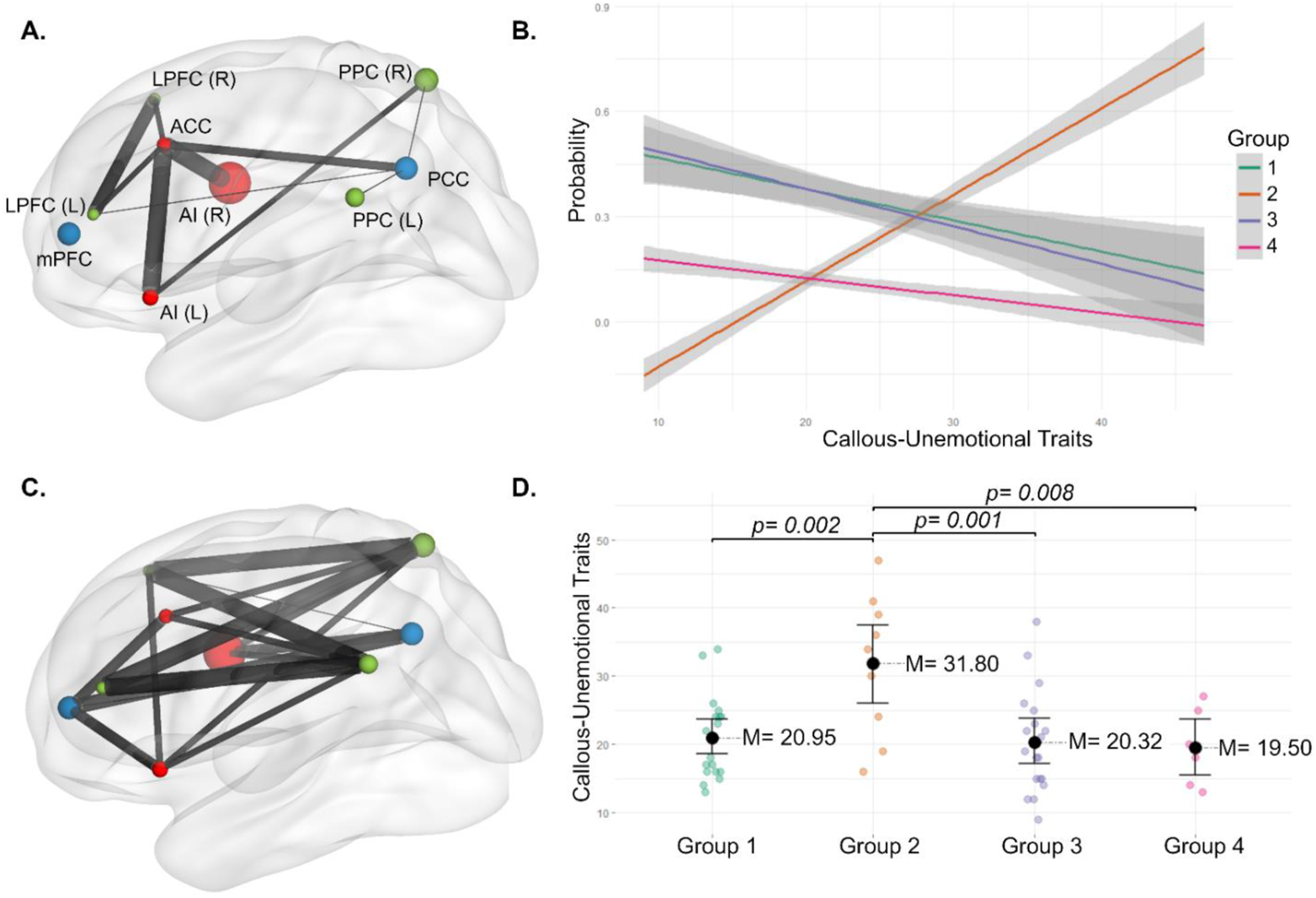
Group-level associations of subgroup two with callous-unemotional traits. A. Depiction of the shared connections for subgroup two. Size of spherical nodes indicate within network centrality (bigger node spheres = more centrality) and size of edges indicate connection density (thicker connection = more density); B. Depicting probability of subgroup inclusion in the presence of total callous-unemotional traits; C. Depicting the heterogeneity of all unshared connections in subgroup 2; D. Mean differences in CU traits by GIMME identified network subgroup. Note: only significant pairwise tests are shown. Node colors: blue = default mode network, red = salience network, green = frontoparietal network.

**Table 1.** Probability of inclusion in GIMME identified subgroups for callous unemotional traits.

|  | Subgroup 2 |  |  |  | Subgroup 3 |  |  |  | Subgroup 4 |  |  |  |
| --- | --- | --- | --- | --- | --- | --- | --- | --- | --- | --- | --- | --- |
| | $\beta$ | S.E. | OR | FDR p | $\beta$ | S.E. | OR | FDR p | $\beta$ | S.E. | OR | FDR p |
| Callous-Unemotional Traits | 0.171* | 0.066 | 1.187 | 0.048 | -0.001 | 0.053 | 0.991 | 0.966 | -0.047 | 0.083 | 0.954 | 0.920 |
| Conduct Issues | -0.105 | 0.094 | 0.900 | 0.438 | -0.215 | 0.086 | 0.801 | 0.064 | 0.009 | 0.092 | 1.009 | 0.920 |
| Tanner Stage | 0.057 | 0.507 | 1.059 | 0.909 | 0.200 | 0.367 | 1.222 | 0.966 | 0.074 | 0.496 | 1.077 | 0.921 |
| Male | 0.461 | 1.024 | 1.584 | 0.816 | -0.031 | 0.751 | 0.969 | 0.966 | -0.885 | 1.043 | 0.412 | 0.920 |

Note: reference category is subgroup 1
\* = FDR corrected $p < 0.05$

#### Analysis confirming subgroup differences on callous-unemotional traits

Subgroups were significantly different on mean level of CU traits and, specifically, the second subgroup had the highest mean level of CU traits (F(3.50)=6.51, *p*= 0.001; Figure 1 Panel C.).

#### Exploratory analysis on dimensional ICU subscale

Uncaring subscale was positively associated with an increase in odds that participants would fall into subgroup two (*β* = 0.195, *p* = 0.048, odds ratio = 1.21, see Supplementary Table 5 and Supplementary Figure 4).

### 3.4. Individual-Specific Features Associated with Callous-Unemotional Traits

#### Individual-specific resting-state network density and callous-unemotional traits

CU traits associated with both within and between network density. Increases in CU traits associated with decreases in number of positive connections within the FPN (*β* = – 0.001, *p _(FDR corrected)_* = 0.042, Figure 2 Panel B and Table 2). For between network density, higher levels of CU traits associated with greater number of positive connections between the DMN-FPN (*β* = 0.002, *p _(FDR corrected)_* = 0.023, Figure 2 Panel C and Table 3).

**Figure 2.**
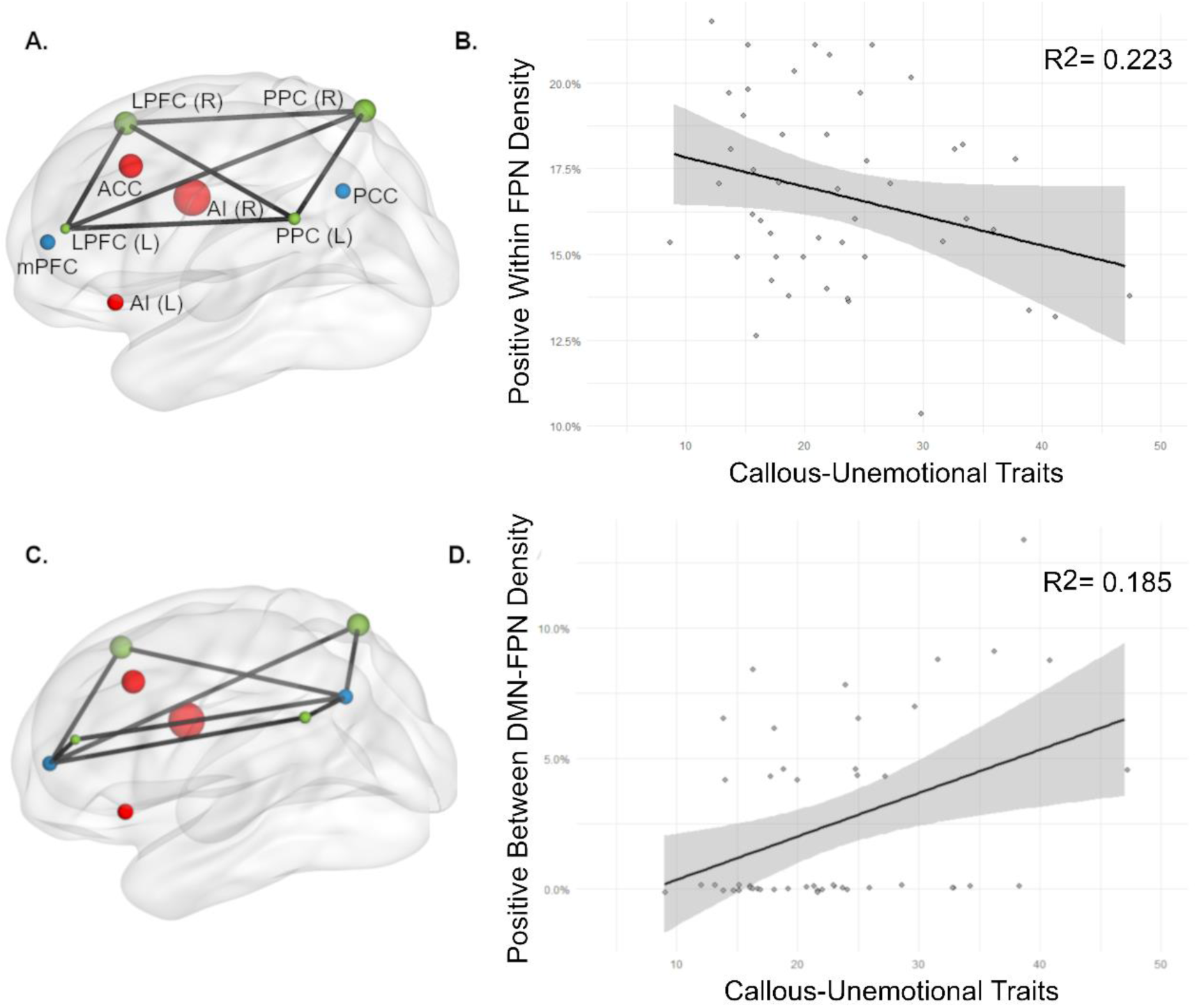
Individual-level associations across all participants with callous-unemotional traits. A. Depiction off all potential connections in the frontoparietal network (FPN); B. Association between positive connections within the FPN; C. Depiction of all possible connections between default mode-frontoparietal networks (DMN-FPN); D. Association between CU traits and positive connections between DMN-FPN. Node: blue = DMN, red = salience network (SAL), green = FPN.

**Table 2.** Within network density across participants

| Network connections | $\beta$ | S.E. | Z | p | FDR p |
| --- | --- | --- | --- | --- | --- |
| **DMN positive** ($R^2 = 0.151$) | | | | | |
| CU traits | -0.001 | 0.001 | -1.217 | 0.224 | 0.447 |
| Tanner | 0.006 | 0.006 | 0.963 | 0.335 | 0.447 |
| Conduct Issues | -0.002 | 0.001 | -2.258 | 0.024 | 0.096 |
| Sex | -0.005 | 0.011 | -0.408 | 0.683 | 0.683 |
| **SAL positive** ($R^2 = 0.098$) | | | | | |
| CU traits | 0.000 | 0.001 | -0.124 | 0.901 | 0.921 |
| Tanner | 0.005 | 0.006 | 0.855 | 0.393 | 0.785 |
| Conduct Issues | -0.002 | 0.001 | -2.098 | 0.036 | 0.144 |
| Sex | 0.001 | 0.012 | 0.099 | 0.921 | 0.921 |
| **SAL negative** ($R^2 = 0.095$) | | | | | |
| CU traits | 0.000 | 0.000 | -0.846 | 0.397 | 0.795 |
| Tanner | 0.000 | 0.003 | 0.122 | 0.903 | 0.903 |
| Conduct Issues | -0.001 | 0.001 | -1.750 | 0.08 | 0.321 |
| Sex | 0.002 | 0.007 | 0.295 | 0.768 | 0.903 |
| **FPN positive** ($R^2 = 0.223$) | | | | | |
| CU traits | -0.001* | 0.000 | -2.557 | 0.011 | 0.042 |
| Tanner | 0.008 | 0.004 | 2.132 | 0.033 | 0.066 |
| Conduct Issues | -0.001 | 0.001 | -1.689 | 0.091 | 0.122 |
| Sex | -0.003 | 0.007 | -0.418 | 0.676 | 0.676 |
| **FPN negative** ($R^2 = 0.136$) | | | | | |
| CU traits | 0.000 | 0.000 | -1.319 | 0.187 | 0.374 |
| Tanner | -0.001 | 0.002 | -0.672 | 0.501 | 0.502 |
| Conduct Issues | -0.001 | 0.000 | -1.556 | 0.12 | 0.374 |
| Sex | 0.004 | 0.004 | 0.862 | 0.389 | 0.502 |

Note: there were no negative DMN connections analysis was not conducted on his outcome.
\*= FDR $p < 0.05$

**Table 3.** Between network density

| Network connections | $\beta$ | S.E. | Z | p | FDR p |
| --- | --- | --- | --- | --- | --- |
| **DMN-FPN positive ($R^2 = 0.185$)** |  |  |  |  |  |
| CU traits | 0.002* | 0.001 | 2.753 | 0.006 | 0.024 |
| Tanner | 0.004 | 0.005 | 0.776 | 0.437 | 0.443 |
| Conduct Issues | -0.001 | 0.001 | -1.143 | 0.253 | 0.443 |
| Sex | -0.007 | 0.010 | -0.767 | 0.443 | 0.443 |
| **DMN-FPN negative ($R^2 = 0.137$)** |  |  |  |  |  |
| CU traits | 0.002 | 0.001 | 2.242 | 0.025 | 0.100 |
| Tanner | 0.006 | 0.006 | 0.918 | 0.359 | 0.718 |
| Conduct Issues | 0.000 | 0.001 | -0.353 | 0.724 | 0.877 |
| Sex | 0.002 | 0.012 | 0.155 | 0.877 | 0.877 |
| **DMN-SAL positive ($R^2 = 0.187$)** |  |  |  |  |  |
| CU traits | 0.001 | 0.001 | 1.847 | 0.065 | 0.129 |
| Tanner | -0.006 | 0.005 | -1.264 | 0.206 | 0.275 |
| Conduct Issues | 0.002 | 0.001 | 2.059 | 0.039 | 0.129 |
| Sex | 0.009 | 0.009 | 0.948 | 0.343 | 0.343 |
| **DMN-SAL negative ($R^2 = 0.202$)** |  |  |  |  |  |
| CU traits | 0.001 | 0.001 | 0.957 | 0.339 | 0.339 |
| Tanner | -0.006 | 0.005 | -1.136 | 0.256 | 0.339 |
| Conduct Issues | 0.002 | 0.001 | 2.473 | 0.013 | 0.054 |
| Sex | 0.018 | 0.010 | 1.759 | 0.079 | 0.157 |
| **SAL-FPN positive ($R^2 = 0.176$)** |  |  |  |  |  |
| CU traits | -0.001 | 0.001 | -1.393 | 0.163 | 0.327 |
| Tanner | -0.005 | 0.006 | -0.759 | 0.448 | 0.448 |
| Conduct Issues | 0.003* | 0.001 | 2.593 | 0.01 | 0.038 |
| Sex | -0.012 | 0.012 | -0.96 | 0.337 | 0.448 |
| **SAL-FPN negative ($R^2 = 0.153$)** |  |  |  |  |  |
| CU traits | -0.001 | 0.001 | -1.484 | 0.138 | 0.184 |
| Tanner | -0.01 | 0.006 | -1.674 | 0.094 | 0.184 |
| Conduct Issues | 0.002 | 0.001 | 1.691 | 0.091 | 0.184 |
| Sex | -0.009 | 0.012 | -0.723 | 0.47 | 0.470 |

\* = FDR $p < 0.05$

#### Individual-specific node centrality and callous-unemotional traits

For within network, CU traits associated with increased positive node centrality in the left insula within the SAL (*β* = 0.010, *p _(FDR corrected)_* = 0.009) and decreases in both positive and negative centrality of the right lateral prefrontal cortex in the FPN (*β* = – 0.005, *p _(FDR corrected)_* = 0.019, *β* = – 0.010, *p _(FDR corrected)_* = 0.006 [respectively], see Supplementary Tables 6-8). For between networks, CU traits associated with central between DMN-FPN connections positively for PCC- left LPFC and negatively for PCC – right PCC (*β* = 0.002, *p _(FDR corrected)_* < 0.001; *β* = 0.004, *p _(FDR corrected)_* = 0.019 [respectively]). Central DMN-SAL network connections showed a positive association between CU traits and positive centrality and a negative association with negative centrality between the PCC-ACC (β = 0.003, *p _(FDR corrected)_* = 0.014; *β* = – 0.002, *p _(FDR corrected)_* = 0.024 [respectively]; Supplementary Tables 9-14, depiction of node centrality Figure 2 Panel A.).

##### Exploratory analyses on individual-specific network features

Facets underlying callous unemotional traits (i.e., callousness, unemotional, and uncaring) had no association with network density features (Supplementary Tables 15-16); but there were associations with centrality network features. The callousness subscale of the ICU associated with decreases in positive network centrality of the right lateral prefrontal cortex within the FPN (*β* = – 0.015, *p* = 0.012) and negatively for positive connections between the posterior cingulate cortex and left lateral prefrontal cortex for between DMN-FPN connectivity (*β* = – 0.005, *p* = 0.009; see Supplementary Figures 17-19). The uncaring subscale negatively associated with positive network centrality of the left insula within the SAL (*β* = 0.013, *p* = 0.033) and negatively with negative network centrality of the right lateral prefrontal cortex within the FPN (*β* = – 0.013, *p* = 0.044). No other significant findings were revealed for ICU subscales (supplementary tables 20-25).

##### Covariate results on individual-specific network features

Covariates also had results of note. Externalizing symptoms positively associated with increases in positive connections between the SAL and FPN (*β* = 0.003, *p _(FDR corrected)_* = 0.038) and negatively associated with both positive and negative network centrality of the anterior cingulate cortex within the salience network (*β* = – 0.012, *p _(FDR corrected)_* = 0.049, *β* = – 0.028, *p _(FDR corrected)_* = 0.034 [respectively]). Pubertal stage positively associated with negative centrality between DMN-SAL at the mPFC-left insula (β = 0.011, *p _(FDR corrected)_* = 0.024) and negative SAL-FPN centrality at the insula(L)-LPFC(R) (β = 0.009, *p _(FDR corrected)_* = 0.003). Interestingly sex had no significant associations after controlling for multiple comparisons.

## 4. Discussion

Results from a community sample of adolescents reveals significant heterogeneity in resting state network connections and supports the notion that CU traits associate with network disintegration. Data-driven results revealed a subset of adolescents with few shared and many individual heterogeneous connections that had higher CU traits, which was independent of conduct problems. This suggests CU traits may lead to more individual-specific alterations in neural circuitry. Individual-specific features indicated less density within the FPN and greater density between the DMN and FPN associated with higher CU traits. Within the FPN this is likely due to fewer connections with the right lateral prefrontal cortex; between DMN-FPN this is likely due to more positive connections between the posterior cingulate cortex-bilateral lateral prefrontal cortices. The present findings connect both adolescent and adult literature on similar features – suggesting 1) psychopathy may be a neurodevelopmental disorder with ontogeny of affective features in youth CU traits and 2) that this can be studied by examining disintegration between the SAL, DMN, and FPN. We extend the current line of research methodologically by examining topological features that modeled both heterogeneity and lagged connections of adolescent brains to reveal the information processing between networks, specifically between DMN-FPN with within the FPN, may be of particular importance.

### 4.1. Network Connection Heterogeneity

The heterogeneity revealed in the present analysis even within identified subgroups emphasizes the issues of relying on the prevalent averaging approaches that hide important individual specific features. For example, the lagged connections modeled by GIMME would have been missed with common network approaches that average across entire timeseries. Because lagged connection are critical features of resting state networks (Beltz & Molenaar, 2015; Mitra et al., 2014) and simulations studies demonstrate GIMME is both accurate and robust (Gates & Molenaar, 2012), the network connections identified here are plausibly more specific to adolescents for which we can make more precise inference on CU traits relation to these network features.

### 4.2. Subgroup Two associates with Greater Callous-Unemotional Traits

The finding that a subgroup had higher representation of CU traits is novel and, if replicated, could provide a base for investigating the heterogeneity of individual connections represented in the group. Exploratory analyses revealed the uncaring CU trait subscale also increased probability in the same subgroup. Although this finding would not have withstood multiple comparison correction, it may be worth further examination to understand whether uncaring drives the association with subgroup two. This subgroup may be replicated and used in future studies to examine shared and heterogeneous connections amongst adolescents with CU traits.

### 4.3. Individual-Specific Network Features Underlie Callous-Unemotional Traits

Individual-specific findings are consistent with theory of dis-integration between networks; and the consistency with the adult literature partially supports the idea that psychopathy is a neurodevelopmental disorder with ontogeny in youth. First, we found that as CU traits increase, the density of connections within the FPN decreases and that the right lateral prefrontal cortex is a central node. This is consistent with adult literature demonstrating interpersonal features of psychopathy negatively associate with node connectivity within the FPN (Philippi et al., 2015), which may indicate a developmental feature of the brain underlying adult psychopathy. Future work examining FPN connections longitudinally could determine this. The FPN is implicated in various aspects of attention alerting, orienting, and cognitive control (e.g., Corbetta, 1998; Scolari et al., 2015; Shulman et al., 2010). Multiple lines of research converge on higher order cognitive processes being impaired in psychopathy (e.g., Baskin-Sommers, Brazil, et al., 2015; Delfin et al., 2018; Maes & Brazil, 2013) and CU traits (e.g., Gluckman et al., 2016; Javakhishvili & Vazsonyi, 2021; Racer et al., 2011). In the context of the literature, the present finding of reduced information flow within the FPN may be related to attention impairments observed in the behavioral literature and it a plausible mechanism for future investigation.

For between network features, our present findings are consistent with both adult findings by Dotterer et al. (2020) and adolescent findings Pu et al. (2017) indicating psychopathic and CU traits associated with greater between DMN-FPN connection density and less anticorrelation (respectively). We extend this work to reveal greater connection density found in adults is present in adolescents with CU traits as well – suggesting a disintegration between networks that may have developmental underpinnings prior to adult psychopathy.

Less connectivity between the DMN-FPN is implicated healthy cognitive functioning, such as social working memory (Xin & Lei, 2015) and cognitive control (Marek et al., 2015; Sheffield et al., 2015), as well as socio-affective processes, such as the interpersonal component of emotional intelligence (Takeuchi et al., 2013) and empathy (Xin & Lei, 2015). Greater connectivity is implicated in aberrant cognitive function (Menon, 2011) and associated with conditions with cognitive and socio-affective impairments such as psychopathy (Dotterer et al., 2020) and schizophrenia (Anticevic et al., 2013; Palaniyappan et al., 2013). Because the FPN suppresses the DMN to improve externally focused task performance (e.g., DeSerisy et al., 2021), these findings suggest cognitive and socio-affective impairments associated with CU traits may be due to failure to suppress the DMN during task directed behavior. Thus, it is plausible that increased information flow between DMN-FPN may interfere with cognitive processes involved in social affective and higher order cognitive functioning. Future studies could examine whether increased DMN-FPN connectivity underlies those with CU traits difficulties in socio-affective tasks or prosocial behavior; and examining the development of between DMN-FPN connections over time may reveal the ontogeny underlying the development of psychopathy.

We did not find aberrant connectivity within the SAL or DMN as hypothesized. However, this was also found in an adult study of psychopathy examining individual network topology (Dotterer et al., 2020). Given that previous studies in adolescents have been inconsistent by not finding aberrant connectivity in the SAL (Umbach & Tottenham, 2020) or DMN (Pu et al., 2017) whereas others have, it is plausible that this may be due differences in modeling approach where using averages across the entire time series are known to create spurious results (Molenaar, 2004). It may also be that the nodes we selected a priori based on previous studies did not include subsystems outside the core regions and, thus, did not demonstrate the disintegration that previous studies have. While it may be argued that adding nodes outside the core nodes for each network may present spurious results that are extraneous to core functions of a network – future studies examining whether the inclusion of subsystems of networks in addition to the core regions may reveal important differences when trying to target specific underlying processes or interactions with subsystems.

### 4.4. Limitations

While the present study presents many strengths there are some limitations to consider. First, this is a small sample size and, thus, may have failed to identify some connections due to a limited power. Also, the results with this community sample may not generalize to forensic samples. However, it is important to note that there is considerable support that CU traits is dimensionally present in the community and demonstrate a range of the same neurocognitive correlates as clinical/forensic samples (Viding & McCrory, 2012). Future studies could build on this research by examining larger samples that include community and forensic samples or oversampling for higher CU traits in community samples.

## 5. Conclusion

The present study demonstrates heterogeneous functional connections in the brains of a community sample of adolescents. The present approach improves upon traditional methods by modeling both contemporaneous, lagged, and directional connections to derive topological features while accounting for individual heterogeneity – improving inference of network features with CU traits. CU traits was more represented in one data-driven identified subgroup that was heterogeneous. Individual-specific features demonstrated lower in density within the FPN and higher density between the DMN and FPN associated with higher CU traits. These findings held whether or not we included conduct problems and withstood multiple comparisons correction. Overall, the present findings suggest less efficiency within the FPN and between DMN-FPN, which are associated with attention, cognitive control, and socio-affective functioning. In the context of the literature on impairments amongst youth with CU traits, these results may explain common impairments amongst these youth. The consistency of these findings with adult literature may suggest neurodevelopmental disorder for adult psychopathy worth further investigation. Disintegration observed in the FPN and between DMN-FPN information processing streams provides support for disintegration between these core networks when examining CU and psychopathic traits. Overall, we demonstrate the importance of modeling individual heterogeneity of network patterns to improve inferences to understand CU traits in adolescents. Modeling individual heterogeneity that investigates topological features of large-scale networks can reveal important neural underpinnings of youth with CU traits that can be used to improve development of individualized treatment approaches for these youth.

## References

Achenbach, T. M., & Rescorla, L. A. (2001). Manual for ASEBA schoolage forms & profiles. University of Vermont, Research Center for Children, Youth, & Families. http://www.aseba.org/

Anticevic, A., Repovs, G., & Barch, D. M. (2013, Jan). Working memory encoding and maintenance deficits in schizophrenia: neural evidence for activation and deactivation abnormalities. Schizophr Bull, 39(1), 168–178. https://doi.org/10.1093/schbul/sbr107

Barry, C. T., Frick, P. J., DeShazo, T. M., McCoy, M., Ellis, M., & Loney, B. R. (2000, May). The importance of callous-unemotional traits for extending the concept of psychopathy to children. J Abnorm Psychol, 109(2), 335–340. https://doi.org/10.1037/0021-843x.109.2.335

Baskin-Sommers, A. R., Brazil, I. A., Ryan, J., Kohlenberg, N. J., Neumann, C. S., & Newman, J. P. (2015). Mapping the association of global executive functioning onto diverse measures of psychopathic traits. Personality Disorders, 6(4), 336–346. https://doi.org/10.1037/per0000125

Baskin-Sommers, A. R., Newman, J. P., Sathasivam, N., & Curtin, J. J. (2011). Evaluating the generalizability of a fear deficit in psychopathic African American offenders. J Abnorm Psychol, 120(1), 71.

Baskin-Sommers, A. R., Waller, R., Fish, A. M., & Hyde, L. W. (2015, Nov). Callous-unemotional traits trajectories interact with earlier conduct problems and exec-utive control to predict violence and substance use among high risk male adolescents. J Abnorm Child Psychol, 43(8), 1529–1541. https://doi.org/10.1007/s10802-015-0041-8

Beltz, A. M. (2018). Gendered mechanisms underlie the relation between pubertal timing and adult depressive symptoms. Journal of Adolescent Health, 62(6), 722–728.

Beltz, A. M., & Gates, K. M. (2017). Network mapping with GIMME. Multivariate Behavioral Research, 52(6), 789–804.

Beltz, A. M., & Molenaar, P. (2015). A posteriori model validation for the temporal order of directed functional connectivity maps. Frontiers in neuroscience, 9, 304.

Benjamini, Y., & Hochberg, Y. (1995). Controlling the false discovery rate: a practical and powerful approach to multiple testing. Journal of the Royal statistical society: series B (Methodological), 57(1), 289–300.

Benjamini, Y., & Yekutieli, D. (2001). The control of the false discovery rate in multiple testing under dependency. Annals of statistics, 1165–1188.

Blair, R. J. (2008). Fine cuts of empathy and the amygdala: dissociable deficits in psychopathy and autism. The Quarterly Journal of Experimental Psychology, 61(1), 157–170.

Blair, R. J. (2010, Feb). Neuroimaging of psychopathy and antisocial behavior: a targeted review. Curr Psychiatry Rep, 12(1), 76–82. https://doi.org/10.1007/s11920-009-0086-x

Blair, R. J., Budhani, S., Colledge, E., & Scott, S. (2005, Mar). Deafness to fear in boys with psychopathic tendencies. J Child Psychol Psychiatry, 46(3), 327–336. https://doi.org/10.1111/j.1469-7610.2004.00356.x

Brown, T. A. (2015). Confirmatory factor analysis for applied research. Guilford publications.

Byrd, A. L., Kahn, R. E., & Pardini, D. A. (2013). A validation of the Inventory of Callous-Unemotional Traits in a community sample of young adult males. Journal of Psychopathology and Behavioral Assessment, 35(1), 20–34.

Caldwell, B. M., Anderson, N. E., Harenski, K. A., Sitney, M. H., Caldwell, M. F., Van Rybroek, G. J., & Kiehl, K. A. (2019). The structural brain correlates of callous-unemotional traits in incarcerated male adolescents. NeuroImage: Clinical, 22, 101703.

Cameron, J. L. (2004). Interrelationships between hormones, behavior, and affect during adolescence: understanding hormonal, physical, and brain changes occurring in association with pubertal activation of the reproductive axis. Introduction to part III. Annals of the New York Academy of Sciences, 1021(1), 110–123.

Cardinale, E. M., O’Connell, K., Robertson, E. L., Meena, L. B., Breeden, A. L., Lozier, L. M., VanMeter, J. W., & Marsh, A. A. (2019). Callous and uncaring traits are associated with reductions in amygdala volume among youths with varying levels of conduct problems. Psychological medicine, 49(9), 1449–1458. https://doi.org/10.1017/S0033291718001927

Cohen, J. R., & D’Esposito, M. (2016). The segregation and integration of distinct brain networks and their relationship to cognition. Journal of Neuroscience, 36(48), 12083–12094.

Cohen, M. A., & Piquero, A. R. (2009). New evidence on the monetary value of saving a high risk youth. Journal of Quantitative Criminology, 25(1), 25–49. https://doi.org/10.1007/s10940-008-9057-3

Cohn, M. D., Pape, L. E., Schmaal, L., Van Den Brink, W., Van Wingen, G., Vermeiren, R. R. J. M., Doreleijers, T. A. H., Veltman, D. J., & Popma, A. (2015). Differential relations between juvenile psychopathic traits and resting state network connectivity. Human Brain Mapping, 36(6), 2396–2405. https://doi.org/10.1002/hbm.22779

Cohn, M. D., Viding, E., McCrory, E., Pape, L., van den Brink, W., Doreleijers, T. A. H., Veltman, D. J., & Popma, A. (2016, 2016/08/30/). Regional grey matter volume and concentration in at-risk adolescents: Untangling associations with callous-unemotional traits and conduct disorder symptoms. Psychiatry Research: Neuroimaging, 254, 180–187. https://doi.org/https://doi.org/10.1016/j.pscychresns.2016.07.003

Corbetta, M. (1998). Frontoparietal cortical networks for directing attention and the eye to visual locations: Identical, independent, or overlapping neural systems? Proceedings of the National Academy of Sciences, 95(3), 831–838. https://doi.org/10.1073/pnas.95.3.831

Dadds, M. R., Jambrak, J., Pasalich, D., Hawes, D. J., & Brennan, J. (2011, Mar). Impaired attention to the eyes of attachment figures and the developmental origins of psychopathy. J Child Psychol Psychiatry, 52(3), 238–245. https://doi.org/10.1111/j.1469-7610.2010.02323.x

Dahl, R. E. (2004). Adolescent brain development: a period of vulnerabilities and opportunities. Keynote address. Annals of the New York Academy of Sciences, 1021(1), 1–22.

De Vico Fallani, F., Richiardi, J., Chavez, M., & Achard, S. (2014, Oct 5). Graph analysis of functional brain networks: practical issues in translational neuroscience. Philos Trans R Soc Lond B Biol Sci, 369(1653). https://doi.org/10.1098/rstb.2013.0521

Delfin, C., Andiné, P., Hofvander, B., Billstedt, E., & Wallinius, M. (2018). Examining Associations Between Psychopathic Traits and Executive Functions in Incarcerated Violent Offenders. Frontiers in Psychiatry, 9, 310–310. https://doi.org/10.3389/fpsyt.2018.00310

DeSerisy, M., Ramphal, B., Pagliaccio, D., Raffanello, E., Tau, G., Marsh, R., Posner, J., & Margolis, A. E. (2021, Apr). Frontoparietal and default mode network connectivity varies with age and intelligence. Dev Cogn Neurosci, 48, 100928. https://doi.org/10.1016/j.dcn.2021.100928

Dotterer, H. L., Hyde, L. W., Shaw, D. S., Rodgers, E. L., Forbes, E. E., & Beltz, A. M. (2020). Connections that characterize callousness: Affective features of psychopathy are associated with personalized patterns of resting-state network connectivity. NeuroImage: Clinical, 28, 102402. https://doi.org/10.1016/j.nicl.2020.102402

Efferson, L. M., & Glenn, A. L. (2018). Examining gender differences in the correlates of psychopathy: A systematic review of emotional, cognitive, and morality-related constructs. Aggression and Violent Behavior, 41, 48–61.

Espinoza, F. A., Vergara, V. M., Reyes, D., Anderson, N. E., Harenski, C. L., Decety, J., Rachakonda, S., Damaraju, E., Rashid, B., & Miller, R. L. (2018). Aberrant functional network connectivity in psychopathy from a large (N= 985) forensic sample. Human Brain Mapping, 39(6), 2624–2634.

Essau, C. A., Sasagawa, S., & Frick, P. J. (2006). Callous-unemotional traits in a community sample of adolescents. Assessment, 13(4), 454–469.

Fanti, K. A., Demetriou, C. A., & Kimonis, E. R. (2013, 2013/07/01). Variants of Callous-Unemotional Conduct Problems in a Community Sample of Adolescents. J Youth Adolesc, 42(7), 964–979. https://doi.org/10.1007/s10964-013-9958-9

Fanti, K. A., Kyranides, M. N., Petridou, M., Demetriou, C. A., & Georgiou, G. (2018, Sep). Neurophysiological markers associated with heterogeneity in conduct problems, callous unemotional traits, and anxiety: Comparing children to young adults. Dev Psychol, 54(9), 1634–1649. https://doi.org/10.1037/dev0000505

Finger, E. C., Marsh, A. A., Mitchell, D. G., Reid, M. E., Sims, C., Budhani, S., Kosson, D. S., Chen, G., Towbin, K. E., & Leibenluft, E. (2008). Abnormal ventromedial prefrontal cortex function in children with psychopathic traits during reversal learning. Archives of General Psychiatry, 65(5), 586–594.

Foster, E. M., Jones, D. E., & Group, C. P. P. R. (2005). The high costs of aggression: Public expenditures resulting from conduct disorder. American journal of public health, 95(10), 1767–1772.

Frick, P. J. (2004). The inventory of callous-unemotional traits. Unpublished rating scale.

Frick, P. J., Ray, J. V., Thornton, L. C., & Kahn, R. E. (2014). Can callous-unemotional traits enhance the understanding, diagnosis, and treatment of serious conduct problems in children and adolescents? A comprehensive review. Psychol Bull, 140(1), 1.

Frick, P. J., & Viding, E. (2009, Fall). Antisocial behavior from a developmental psychopathology perspective. Dev Psychopathol, 21(4), 1111–1131. https://doi.org/10.1017/s0954579409990071

Frick, P. J., & White, S. F. (2008). Research review: The importance of callous-unemotional traits for developmental models of aggressive and antisocial behavior. Journal of child psychology and psychiatry, 49(4), 359–375.

Gates, K. M., Lane, S. T., Varangis, E., Giovanello, K., & Guiskewicz, K. (2017, 2017/03/04). Unsupervised Classification During Time-Series Model Building. Multivariate Behavioral Research, 52(2), 129–148. https://doi.org/10.1080/00273171.2016.1256187

Gates, K. M., & Molenaar, P. C. (2012). Group search algorithm recovers effective connectivity maps for individuals in homogeneous and heterogeneous samples. NeuroImage, 63(1), 310–319.

Gates, K. M., Molenaar, P. C., Hillary, F. G., Ram, N., & Rovine, M. J. (2010). Automatic search for fMRI connectivity mapping: an alternative to Granger causality testing using formal equivalences among SEM path modeling, VAR, and unified SEM. NeuroImage, 50(3), 1118–1125.

Gates, K. M., Molenaar, P. C., Hillary, F. G., & Slobounov, S. (2011). Extended unified SEM approach for modeling event-related fMRI data. NeuroImage, 54(2), 1151–1158.

Gluckman, N. S., Hawes, D. J., & Russell, A. M. T. (2016, 2016/08/01). Are Callous-Unemotional Traits Associated with Conflict Adaptation in Childhood? Child Psychiatry & Human Development, 47(4), 583–592. https://doi.org/10.1007/s10578-015-0593-4

Goetschius, L. G., Hein, T. C., McLanahan, S. S., Brooks-Gunn, J., McLoyd, V. C., Dotterer, H. L., Lopez-Duran, N., Mitchell, C., Hyde, L. W., & Monk, C. S. (2020). Association of childhood violence exposure with adolescent neural network density. JAMA network open, 3(9), e2017850–e2017850.

Hadjicharalambous, M. Z., & Fanti, K. A. (2018, Jun). Self Regulation, Cognitive Capacity and Risk Taking: Investigating Heterogeneity Among Adolescents with Callous-Unemotional Traits. Child Psychiatry Hum Dev, 49(3), 331–340. https://doi.org/10.1007/s10578-017-0753-9

Hamilton, R. K., Hiatt Racer, K., & Newman, J. P. (2015, Oct). Impaired integration in psychopathy: A unified theory of psychopathic dysfunction. Psychol Rev, 122(4), 770–791. https://doi.org/10.1037/a0039703

Hawes, D., & Dadds, M. R. (2012). Revisiting the role of empathy in childhood pathways to antisocial behavior.

Herpers, P. C., Scheepers, F. E., Bons, D. M., Buitelaar, J. K., & Rommelse, N. N. (2014). The cognitive and neural correlates of psychopathy and especially callous–unemotional traits in youths: A systematic review of the evidence. Development and psychopathology, 26(1), 245–273.

Hyde, L. W., Burt, S. A., Shaw, D. S., Donnellan, M. B., & Forbes, E. E. (2015). Early starting, aggressive, and/or callous–unemotional? Examining the overlap and predictive utility of antisocial behavior subtypes. J Abnorm Psychol, 124(2), 329–342. https://doi.org/10.1037/abn0000029

Hyde, L. W., Shaw, D. S., Murray, L., Gard, A., Hariri, A. R., & Forbes, E. E. (2016, May). Dissecting the role of amygdala reactivity in antisocial behavior in a sample of young, low-income, urban men. Clin Psychol Sci, 4(3), 527–544. https://doi.org/10.1177/2167702615614511

Insel, T. R. (2014). The NIMH research domain criteria (RDoC) project: precision medicine for psychiatry. American Journal of Psychiatry, 171(4), 395–397.

Javakhishvili, M., & Vazsonyi, A. T. (2021, 2021/02/12). Empathy, Self-control, Callous-Unemotionality, and Delinquency: Unique and Shared Developmental Antecedents. Child Psychiatry & Human Development. https://doi.org/10.1007/s10578-021-01137-2

Jones, A. P., Laurens, K. R., Herba, C. M., Barker, G. J., & Viding, E. (2009, Jan). Amygdala hypoactivity to fearful faces in boys with conduct problems and callous-unemotional traits. Am J Psychiatry, 166(1), 95–102. https://doi.org/10.1176/appi.ajp.2008.07071050

Kaiser, M. (2011, 2011/08/01/). A tutorial in connectome analysis: Topological and spatial features of brain networks. NeuroImage, 57(3), 892–907. https://doi.org/https://doi.org/10.1016/j.neuroimage.2011.05.025

Kiehl, K. A. (2006). A cognitive neuroscience perspective on psychopathy: evidence for paralimbic system dysfunction. Psychiatry Research, 142(2-3), 107–128. https://doi.org/10.1016/j.psychres.2005.09.013

Kimonis, E. R., Frick, P. J., Skeem, J. L., Marsee, M. A., Cruise, K., Munoz, L. C., Aucoin, K. J., & Morris, A. S. (2008). Assessing callous–unemotional traits in adolescent offenders: Validation of the Inventory of Callous–Unemotional Traits. International journal of law and psychiatry, 31(3), 241–252.

Korponay, C., Pujara, M., Deming, P., Philippi, C., Decety, J., Kosson, D. S., Kiehl, K. A., & Koenigs, M. (2017). Impulsive-antisocial dimension of psychopathy linked to enlargement and abnormal functional connectivity of the striatum. Biological Psychiatry: Cognitive Neuroscience and Neuroimaging, 2(2), 149–157.

Lane, S., Gates, K., Fisher, Z., Arizmendi, C., Molenaar, P., Hallquist, M., Pike, H., Henry, T., Duffy, K., Luo, L., Beltz, A. M., Wright, A., Park, J., & Alvarez, S. C. (2021). gimme: Group Iterative Multiple Model Estimation R package version 0.7-5. CRAN. https://doi.org/https://CRAN.R-project.org/package=gimme

Larson, C. L., Baskin-Sommers, A. R., Stout, D. M., Balderston, N. L., Curtin, J. J., Schultz, D. H., Kiehl, K. A., & Newman, J. P. (2013). The interplay of attention and emotion: top-down attention modulates amygdala activation in psychopathy. Cognitive, Affective, & Behavioral Neuroscience, 13(4), 757–770.

Lozier, L. M., Cardinale, E. M., VanMeter, J. W., & Marsh, A. A. (2014, Jun). Mediation of the relationship between callous-unemotional traits and proactive aggression by amygdala response to fear among children with conduct problems. JAMA Psychiatry, 71(6), 627–636. https://doi.org/10.1001/jamapsychiatry.2013.4540

Maes, J. H., & Brazil, I. A. (2013). No clear evidence for a positive association between the interpersonal-affective aspects of psychopathy and executive functioning. Psychiatry Research, 210(3), 1265–1274.

Marek, S., Hwang, K., Foran, W., Hallquist, M. N., & Luna, B. (2015). The Contribution of Network Organization and Integration to the Development of Cognitive Control. PLoS biology, 13(12), e1002328–e1002328. https://doi.org/10.1371/journal.pbio.1002328

Marsh, A. A., Finger, E. C., Fowler, K. A., Adalio, C. J., Jurkowitz, I. T., Schechter, J. C., Pine, D. S., Decety, J., & Blair, R. J. R. (2013). Empathic responsiveness in amygdala and anterior cingulate cortex in youths with psychopathic traits. Journal of child psychology and psychiatry, 54(8), 900–910.

Mayeux, R. (2004). Biomarkers: potential uses and limitations. NeuroRx : the journal of the American Society for Experimental NeuroTherapeutics, 1(2), 182–188. https://doi.org/10.1602/neurorx.1.2.182

Menon, V. (2011). Large-scale brain networks and psychopathology: a unifying triple network model. Trends in cognitive sciences, 15(10), 483–506.

Menon, V. (2015). Salience network. In A. W. Toga (Ed.), Brain Mapping: An Encyclopedic Reference (Vol. 2, pp. 597–611). Elsevier.

Menon, V., & Uddin, L. Q. (2010, 05/29). Saliency, switching, attention and control: a network model of insula function. Brain Structure & Function, 214(5-6), 655–667. https://doi.org/10.1007/s00429-010-0262-0

Mitra, A., Snyder, A. Z., Hacker, C. D., & Raichle, M. E. (2014). Lag structure in resting-state fMRI. Journal of neurophysiology, 111(11), 2374–2391. https://doi.org/10.1152/jn.00804.2013

Molenaar, P. C. M. (2004, 2004/10/01). A Manifesto on Psychology as Idiographic Science: Bringing the Person Back Into Scientific Psychology, This Time Forever. Measurement: Interdisciplinary Research and Perspectives, 2(4), 201–218. https://doi.org/10.1207/s15366359mea0204_1

Neumann, C. S., & Hare, R. D. (2008). Psychopathic traits in a large community sample: links to violence, alcohol use, and intelligence. Journal of consulting and clinical psychology, 76(5), 893.

Nooner, K. B., Colcombe, S. J., Tobe, R. H., Mennes, M., Benedict, M. M., Moreno, A. L., & … Milham, M. P. (2012). The NKI-Rockland Sample: A Model for Accelerating the Pace of Discovery Science in Psychiatry. Frontiers in Neuroscience, 6(152).

Palaniyappan, L., Simmonite, M., White, T. P., Liddle, E. B., & Liddle, P. F. (2013). Neural primacy of the salience processing system in schizophrenia. Neuron, 79(4), 814–828. https://doi.org/10.1016/j.neuron.2013.06.027

Parent, A.-S., Teilmann, G., Juul, A., Skakkebaek, N. E., Toppari, J., & Bourguignon, J.-P. (2003). The Timing of Normal Puberty and the Age Limits of Sexual Precocity: Variations around the World, Secular Trends, and Changes after Migration. Endocrine Reviews, 24(5), 668–693. https://doi.org/10.1210/er.2002-0019

Penny, W. D., Friston, K. J., Ashburner, J. T., Kiebel, S. J., & Nichols, T. E. (2011). Statistical parametric mapping: the analysis of functional brain images. Elsevier.

Perez, V. B., Swerdlow, N. R., Braff, D. L., Näätänen, R., & Light, G. A. (2014). Using biomarkers to inform diagnosis, guide treatments and track response to interventions in psychotic illnesses. Biomarkers in medicine, 8(1), 9–14. https://doi.org/10.2217/bmm.13.133

Petersen, A. C., Crockett, L., Richards, M., & Boxer, A. (1988). A self-report measure of pubertal status: Reliability, validity, and initial norms. J Youth Adolesc, 17(2), 117–133.

Philippi, C. L., Pujara, M. S., Motzkin, J. C., Newman, J., Kiehl, K. A., & Koenigs, M. (2015). Altered resting-state functional connectivity in cortical networks in psychopathy. The Journal of neuroscience : the official journal of the Society for Neuroscience, 35(15), 6068–6078. https://doi.org/10.1523/JNEUROSCI.5010-14.2015

Pons, P., & Latapy, M. (2005). Computing communities in large networks using random walks. International symposium on computer and information sciences,

Price, R. B., Gates, K., Kraynak, T. E., Thase, M. E., & Siegle, G. J. (2017). Data-driven subgroups in depression derived from directed functional connectivity paths at rest. Neuropsychopharmacology, 42(13), 2623–2632.

Pu, W., Luo, Q., Jiang, Y., Gao, Y., Ming, Q., & Yao, S. (2017, Sep 12). Alterations of Brain Functional Architecture Associated with Psychopathic Traits in Male Adolescents with Conduct Disorder. Sci Rep, 7(1), 11349. https://doi.org/10.1038/s41598-017-11775-z

Pujol, J., Batalla, I., Contreras-Rodríguez, O., Harrison, B. J., Pera, V., Hernández-Ribas, R., Real, E., Bosa, L., Soriano-Mas, C., Deus, J., López-Solà, M., Pifarré, J., Menchón, J. M., & Cardoner, N. (2012). Breakdown in the brain network subserving moral judgment in criminal psychopathy. Social Cognitive and Affective Neuroscience, 7(8), 917–923. https://doi.org/10.1093/scan/nsr075

R Core Team. (2021). R: A language and environment for statistical computing. URL https://www.R-project.org/.

Racer, K. H., Gilbert, T. T., Luu, P., Felver-Gant, J., Abdullaev, Y., & Dishion, T. J. (2011). Attention network performance and psychopathic symptoms in early adolescence: an ERP study. Journal of abnormal child psychology, 39(7), 1001–1012. https://doi.org/10.1007/s10802-011-9522-6

Raschle, N. M., Menks, W. M., Fehlbaum, L. V., Steppan, M., Smaragdi, A., Gonzalez-Madruga, K., Rogers, J., Clanton, R., Kohls, G., Martinelli, A., Bernhard, A., Konrad, K., Herpertz-Dahlmann, B., Freitag, C. M., Fairchild, G., De Brito, S. A., & Stadler, C. (2018, 2018/01/01/). Callous-unemotional traits and brain structure: Sex-specific effects in anterior insula of typically-developing youths. NeuroImage: Clinical, 17, 856–864. https://doi.org/https://doi.org/10.1016/j.nicl.2017.12.015

Rezaeinia, P., Fairley, K., Pal, P., Meyer, F. G., & Carter, R. M. (2020). Identifying brain network topology changes in task processes and psychiatric disorders. Network Neuroscience, 4(1), 257–273.

Rogers, J. C., & De Brito, S. A. (2016). Cortical and subcortical gray matter volume in youths with conduct problems: a meta-analysis. JAMA Psychiatry, 73(1), 64–72.

Rosseel, Y. (2012). Lavaan: An R Package for Structural Equation Modeling. Journal of Statistical Software, 48(2), 1–36.

Satterthwaite, T. D., Elliott, M. A., Gerraty, R. T., Ruparel, K., Loughead, J., Calkins, M. E., Eickhoff, S. B., Hakonarson, H., Gur, R. C., & Gur, R. E. (2013). An improved framework for confound regression and filtering for control of motion artifact in the preprocessing of resting-state functional connectivity data. NeuroImage, 64, 240–256.

Scolari, M., Seidl-Rathkopf, K. N., & Kastner, S. (2015). Functions of the human frontoparietal attention network: Evidence from neuroimaging. Current Opinion in Behavioral Sciences, 1, 32–39.

Sebastian, C. L., De Brito, S. A., McCrory, E. J., Hyde, Z. H., Lockwood, P. L., Cecil, C. A., & Viding, E. (2016). Grey matter volumes in children with conduct problems and varying levels of callous-unemotional traits. Journal of abnormal child psychology, 44(4), 639–649.

Sebastian, C. L., McCrory, E. J., Cecil, C. A., Lockwood, P. L., De Brito, S. A., Fontaine, N. M., & Viding, E. (2012). Neural responses to affective and cognitive theory of mind in children with conduct problems and varying levels of callous-unemotional traits. Archives of General Psychiatry, 69(8), 814–822.

Sebastian, C. L., McCrory, E. J. P., Cecil, C. A. M., Lockwood, P. L., De Brito, S. A., Fontaine, N. M. G., & Viding, E. (2012). Neural Responses to Affective and Cognitive Theory of Mind in Children With Conduct Problems and Varying Levels of Callous-Unemotional Traits. Archives of General Psychiatry, 69(8), 814. https://doi.org/10.1001/archgenpsychiatry.2011.2070

Sethi, A., Sarkar, S., Dell’Acqua, F., Viding, E., Catani, M., Murphy, D. G. M., & Craig, M. C. (2018, 2018/04/01/). Anatomy of the dorsal default-mode network in conduct disorder: Association with callous-unemotional traits. Developmental cognitive neuroscience, 30, 87–92. https://doi.org/https://doi.org/10.1016/j.dcn.2018.01.004

Sheffield, J. M., Repovs, G., Harms, M. P., Carter, C. S., Gold, J. M., MacDonald, A. W., 3rd, Daniel Ragland, J., Silverstein, S. M., Godwin, D., & Barch, D. M. (2015). Fronto-parietal and cingulo-opercular network integrity and cognition in health and schizophrenia. Neuropsychologia, 73, 82–93. https://doi.org/10.1016/j.neuropsychologia.2015.05.006

Shulman, G. L., Pope, D. L., Astafiev, S. V., McAvoy, M. P., Snyder, A. Z., & Corbetta, M. (2010). Right hemisphere dominance during spatial selective attention and target detection occurs outside the dorsal frontoparietal network. Journal of Neuroscience, 30(10), 3640–3651.

Sisk, C. L., & Foster, D. L. (2004). The neural basis of puberty and adolescence. Nature Neuroscience, 7(10), 1040–1047.

Smith, S. M., Miller, K. L., Salimi-Khorshidi, G., Webster, M., Beckmann, C. F., Nichols, T. E., Ramsey, J. D., & Woolrich, M. W. (2011). Network modelling methods for FMRI. NeuroImage, 54(2), 875–891.

Sörbom, D. (1989, 1989/09/01). Model modification. Psychometrika, 54(3), 371–384. https://doi.org/10.1007/BF02294623

Stiso, J., & Bassett, D. S. (2018). Spatial embedding imposes constraints on neuronal network architectures. Trends in cognitive sciences, 22(12), 1127–1142.

Takeuchi, H., Taki, Y., Nouchi, R., Sekiguchi, A., Hashizume, H., Sassa, Y., Kotozaki, Y., Miyauchi, C. M., Yokoyama, R., & Iizuka, K. (2013). Resting state functional connectivity associated with trait emotional intelligence. NeuroImage, 83, 318–328.

Uddin, L. Q., Kelly, A. M., Biswal, B. B., Castellanos, F. X., & Milham, M. P. (2009). Functional connectivity of default mode network components: correlation, anticorrelation, and causality. Human Brain Mapping, 30(2), 625–637. https://doi.org/10.1002/hbm.20531

Umbach, R. H., & Tottenham, N. (2020). Callous-unemotional traits and reduced default mode network connectivity within a community sample of children. Development and psychopathology, 1–14.

van der Stouwe, T., Asscher, J. J., Stams, G. J. J. M., Deković, M., & van der Laan, P. H. (2014, 2014/08/01/). The effectiveness of Multisystemic Therapy (MST): A meta-analysis. Clinical Psychology Review, 34(6), 468–481. https://doi.org/https://doi.org/10.1016/j.cpr.2014.06.006

Veroude, K., von Rhein, D., Chauvin, R. J., van Dongen, E. V., Mennes, M. J., Franke, B., Heslenfeld, D. J., Oosterlaan, J., Hartman, C. A., & Hoekstra, P. J. (2016). The link between callous-unemotional traits and neural mechanisms of reward processing: An fMRI study. Psychiatry Research: Neuroimaging, 255, 75–80.

Viding, E., & McCrory, E. J. (2012, Aug). Genetic and neurocognitive contributions to the development of psychopathy. Dev Psychopathol, 24(3), 969–983. https://doi.org/10.1017/s095457941200048x

Viding, E., Sebastian, C. L., Dadds, M. R., Lockwood, P. L., Cecil, C. A., De Brito, S. A., & McCrory, E. J. (2012). Amygdala response to preattentive masked fear in children with conduct problems: the role of callous-unemotional traits. American Journal of Psychiatry, 169(10), 1109–1116.

Wallace, G. L., White, S. F., Robustelli, B., Sinclair, S., Hwang, S., Martin, A., & Blair, R. J. R. (2014). Cortical and subcortical abnormalities in youths with conduct disorder and elevated callous-unemotional traits. Journal of the American Academy of Child & Adolescent Psychiatry, 53(4), 456–465. e451.

Wechsler, D. (2011). Wechsler abbreviated scale of intelligence-(WASI-II) (Vol. 4). NCS Pearson.

White, S. F., Williams, W. C., Brislin, S. J., Sinclair, S., Blair, K. S., Fowler, K. A., Pine, D. S., Pope, K., & Blair, R. J. (2012). Reduced activity within the dorsal endogenous orienting of attention network to fearful expressions in youth with disruptive behavior disorders and psychopathic traits. Development and psychopathology, 24(3), 1105–1116. https://doi.org/10.1017/S0954579412000569

Whitfield-Gabrieli, S., & Nieto-Castanon, A. (2012). Conn: a functional connectivity toolbox for correlated and anticorrelated brain networks. Brain Connect, 2(3), 125–141. https://doi.org/10.1089/brain.2012.0073

Xin, F., & Lei, X. (2015). Competition between frontoparietal control and default networks supports social working memory and empathy. Social Cognitive and Affective Neuroscience, 10(8), 1144–1152.

Yoder, K. J., Lahey, B. B., & Decety, J. (2016). Callous traits in children with and without conduct problems predict reduced connectivity when viewing harm to others. Scientific Reports, 6(1), 20216. https://doi.org/10.1038/srep20216

