## Supplementary material for "Resting-state network topology characterizing callous-unemotional traits in adolescence": supplmentry material

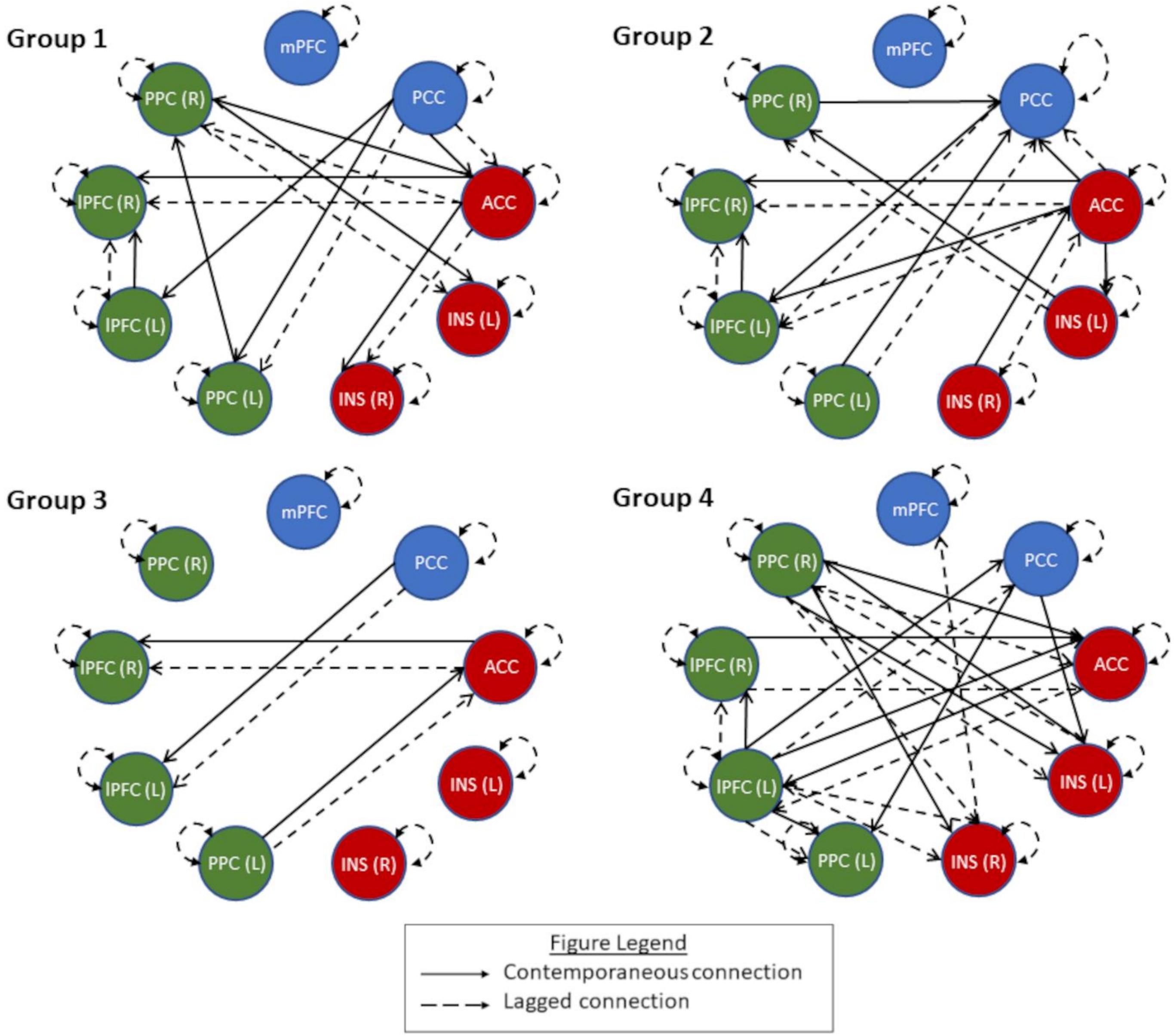


Supplementary Figure 1. Final GIMME identified group network patterns. See supplementary table 3 as well as supplementary figure3 for group other network features.

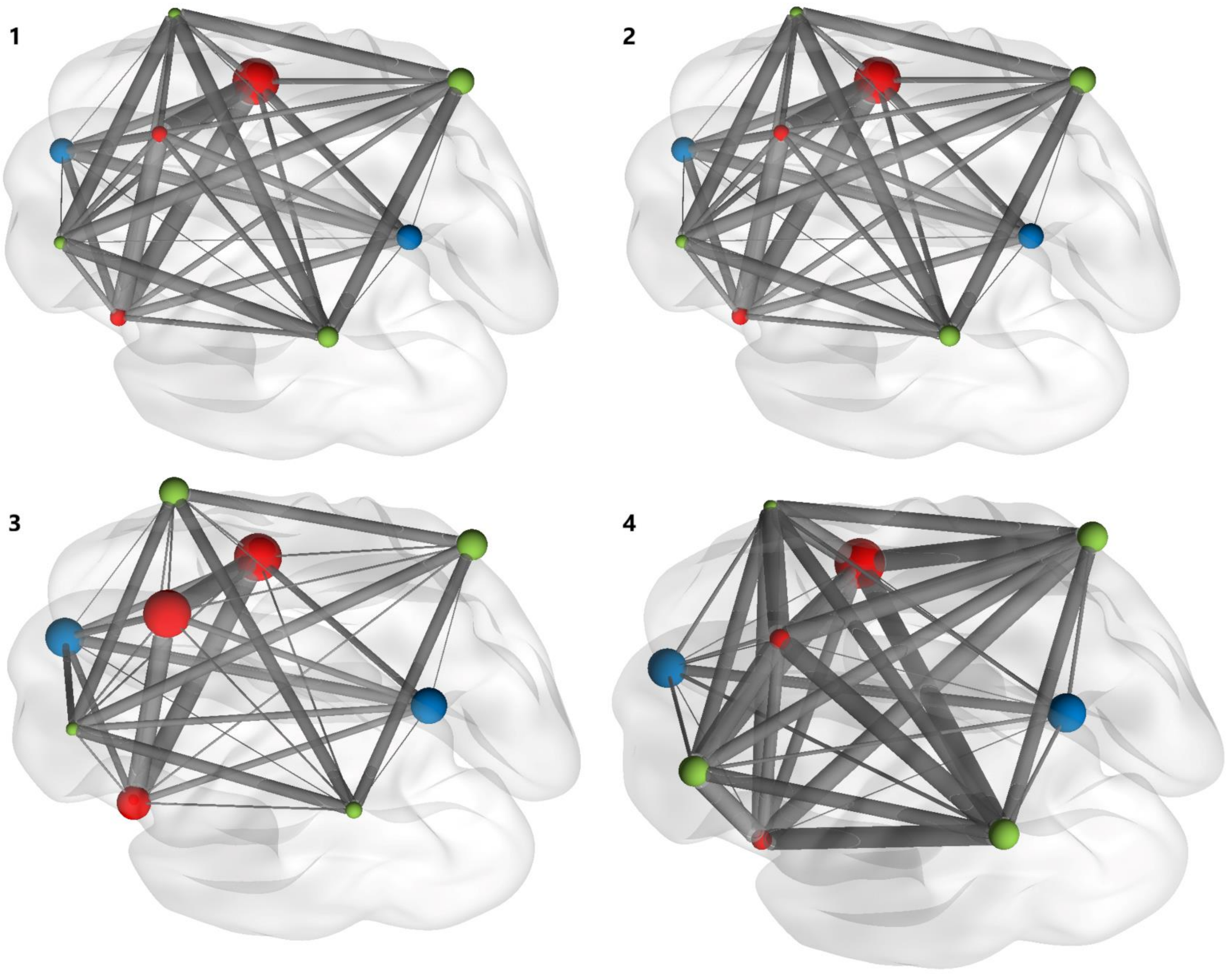


Supplementary Figure 2. Depiction of subgroup mean connections for both shared and unshared connections and node centrality for all participants in each of the four identified groups. Node size indicates within network centrality and connection size indicate connection density. 1= group one, 2=group two, 3 = group three, 4= group four.

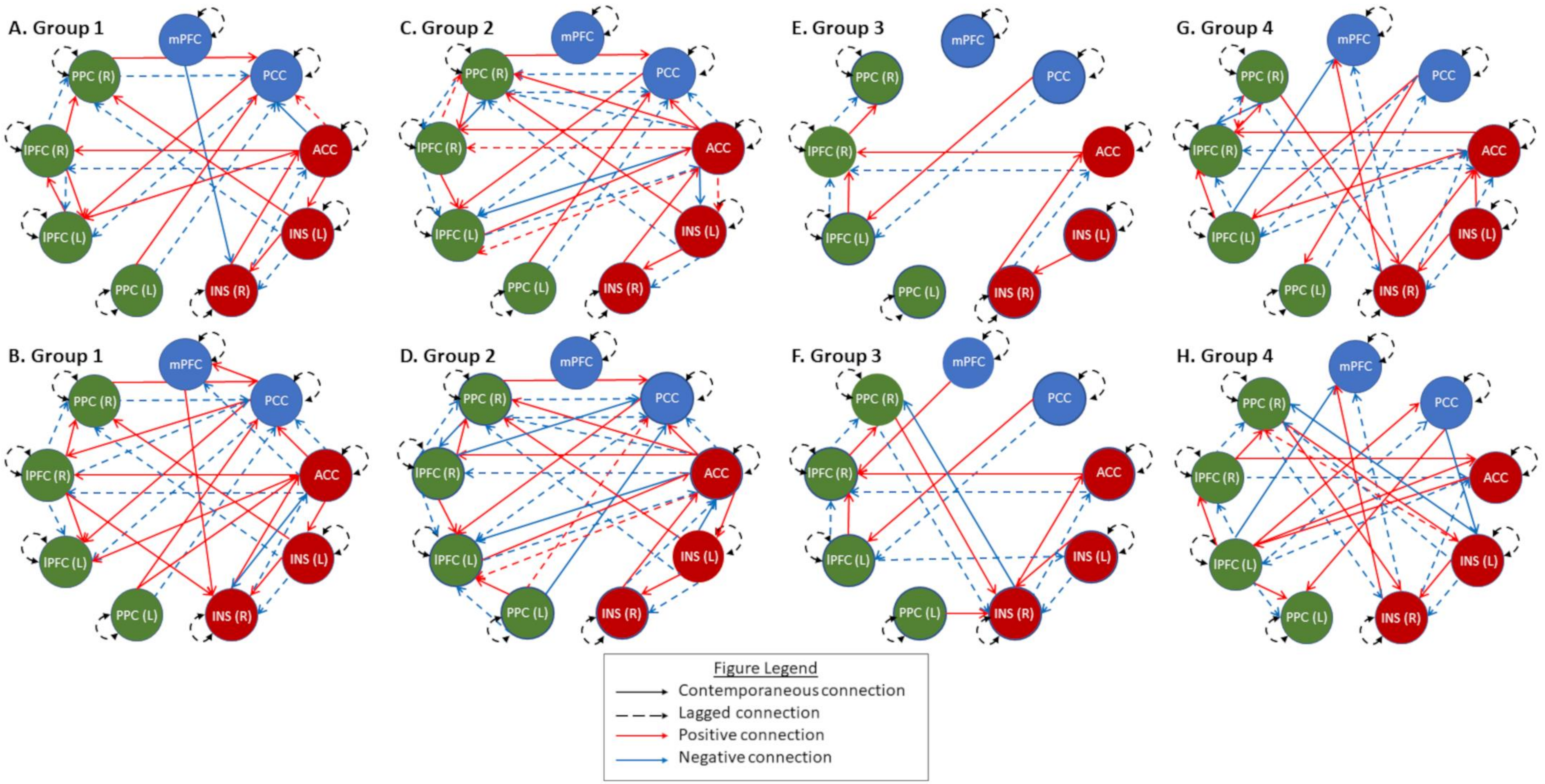


Supplementary Figure 3. Final GIMME networks for eight example participants – two for each group. Solid lines depict contemporaneous connections and dotted lines depict lagged connections with red line indicating positive connections (indicated by a positive β) and blue lines indicating negative connections (indicated by a negative β). Node colors indicate specific networks: blue indicates default mode network, red indicate the salience network, and green regions indicate the frontoparietal network.

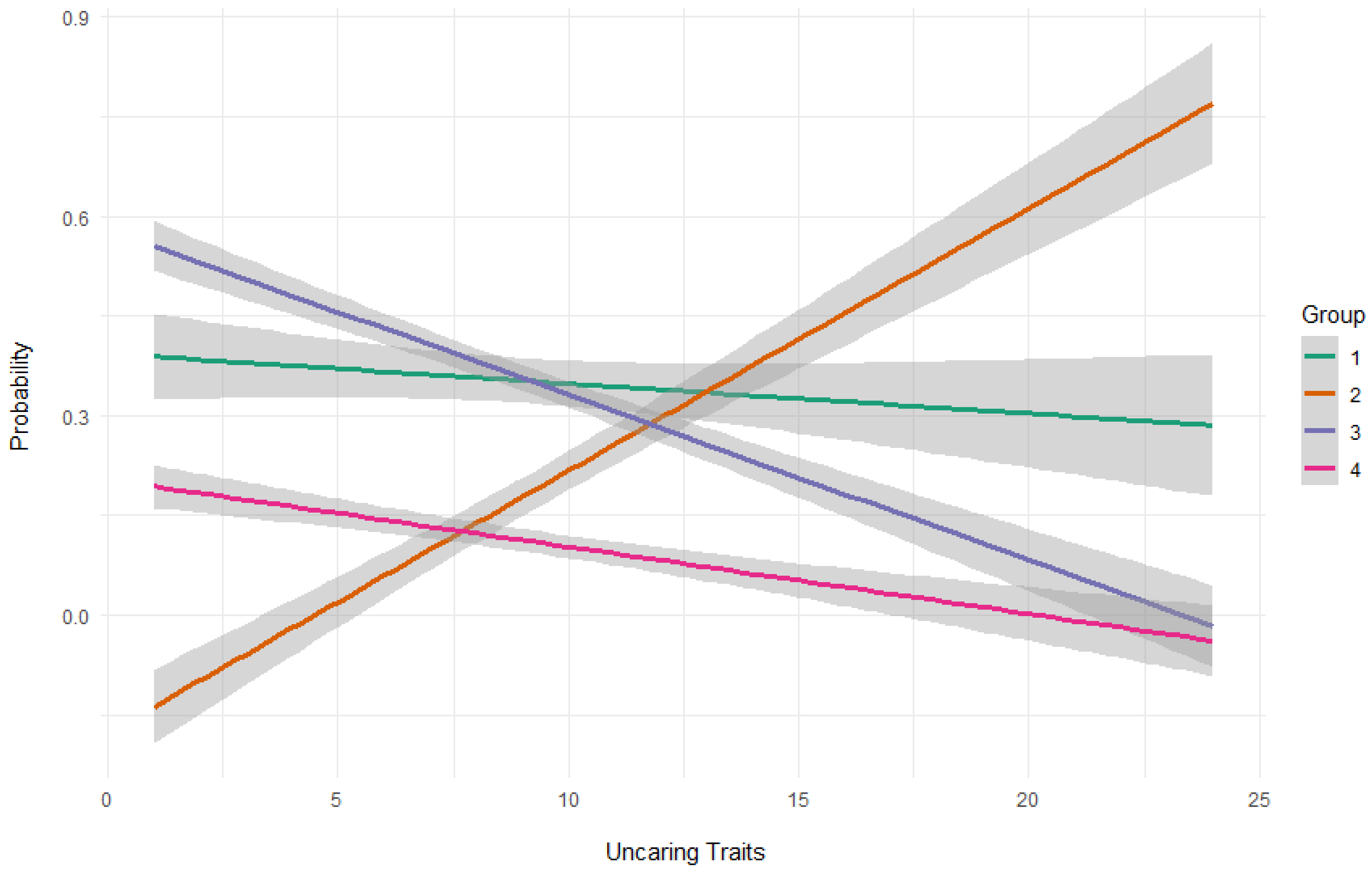


Supplementary Figure 2. Depicting probability of group inclusion because of the uncaring subscale.

Supplementary Table 1. MNI Coordinates for ROIs

| **Network**<br>Region in network | **MNI coordinates**<br>(x, y, z) |
|---|---|
| **Default Mode Network** | |
| Medial Prefrontal Cortex | 1, 55, -3 |
| Posterior Cingulate Cortex | 1, -61, 38 |
| **Salience Network** | |
| Anterior Cingulate Cortex | 0, 22, 35 |
| Anterior Insula (L) | -44, 13, 1 |
| Anterior Insula (R) | 47, 14, 0 |
| **Frontoparietal Network** | |
| Lateral Prefrontal Cortex (L) | -43, 33, 28 |
| Lateral Prefrontal Cortex (R) | 41, 38, 30 |
| Posterior Parietal Cortex (L) | -46, -58, 49 |
| Posterior Parietal Cortex (R) | 52, -52, 45 |

Note: (L) = left, (R) = right

Supplementary Table 2. Model Fit statistics

| Model | Modularity | RMSEA (min-max) | SRMR (min-max) | CFI (min-max) | NFI (min-max) | Met Criteria [a] |
|---|---|---|---|---|---|---|
| Individual | 0.024 | 0.122 (0.160-0.136) | 0.024**+** (0.018-0.033) | 0.953**+** (0.949-0.966) | 0.910 (0.902-0.935) | Yes |
| Group 1 | 0.201 | 0.125 (0.118-0.134) | 0.026**+** (0.020-0.033) | 0.950**+** (0.950-0.954) | 0.907 (0.904-0.915) | Yes |
| Group 2 | 0.200 | 0.126 (0.117-0.136) | 0.024**+** (0.019-0.028) | 0.951**+** (0.950-0.956) | 0.908 (0.902-0.918) | Yes |
| Group 3 | 0.252 | 0.122 (0.106-0.125) | 0.024**+** (0.018-0.028) | 0.950**+** (0.949-0.957) | 0.908 (0.911-0.930) | Yes |
| Group 4 | 0.213 | 0.114 (0.109-0.120) | 0.024**+** (0.019-0.026) | 0.961**+** (0.957-0.966) | 0.925 (0.917-0.935) | Yes |

[a] excellent fit criteria by Brown (2015) - requires two out of four alternative fit indices are met: root mean squared error of approximation (RMSEA)≤0.05, standardized root mean residual (SRMR)≤0.05, comparative fit index (CFI)≥0.95, or non-normed fit index (NNFI)≥0.95

+ = Met fit indices criteria

Supplementary Table 3. Group network density

| Group | Average Connections (sd) | Average Contemporaneous (sd) | Average Lagged (sd) |
|---|---|---|---|
| Group 1 | 62.33 (24.23) | 12.68 (10.87) | 12.19 (12.19) |
| Group 2 | 39.89 (23.54) | 10.75 (9.23) | 19.81 (10.79) |
| Group 3 | 53.67 (25.88) | 10.56 (11.19) | 28.69 (14.65) |
| Group 4 | 20.56 (6.82) | 4.06 (4.55) | 5.91 (5.91) |

Supplementary Table 5. Exploratory Probability of inclusion in GIMME identified groups for callous unemotional trait facets (ICU subscales)

| | Group 2 | | | | Group 3 | | | | Group 4 | | | |
|---|---|---|---|---|---|---|---|---|---|---|---|---|
| | β | S.E. | OR | p | β | S.E. | OR | p | β | S.E. | OR | p |
| Callousness | 0.194 | 0.145 | 1.214 | 0.264 | 0.062 | 0.160 | 1.064 | 0.694 | 0.218 | 0.234 | 1.243 | 0.351 |
| Uncaring | 0.191* | 0.097 | 1.211 | 0.048 | -0.136 | 0.105 | 0.872 | 0.195 | -0.231 | 0.179 | 0.794 | 0.197 |
| Unemotional | 0.113 | 0.203 | 1.119 | 0.577 | 0.159 | 0.141 | 1.173 | 0.260 | 0.062 | 0.198 | 1.063 | 0.754 |
| Tanner Stage | 0.034 | 0.070 | 1.034 | 0.944 | 0.116 | 0.356 | 1.123 | 0.743 | 0.131 | 0.540 | 1.139 | 0.809 |
| Male | 0.487 | 1.007 | 1.628 | 0.590 | 0.181 | 0.718 | 1.199 | 0.800 | -0.569 | 1.058 | 0.565 | 0.628 |

Note: reference category is group 1, *= uncorrected $p < 0.05$

Supplementary Table 6. Within default mode network centrality

| Network connections | β | S.E. | Z | p | FDR p |
|---|---|---|---|---|---|
| **mPFC positive** ($R^2$ = 0.0.07) | | | | | |
| CU traits | 0.003 | 0.002 | 1.129 | 0.259 | 0.442 |
| Tanner | -0.021 | 0.021 | -0.970 | 0.332 | 0.443 |
| Conduct Issues | -0.001 | 0.004 | -0.318 | 0.751 | 0.750 |
| Sex | -0.042 | 0.043 | -0.983 | 0.326 | 0.751 |
| **mPFC negative** ($R^2$ = 0.042) | | | | | |
| CU traits | 0.001 | 0.001 | 0.906 | 0.430 | 0.730 |
| Tanner | -0.006 | 0.005 | -1.141 | 0.263 | 0.730 |
| Conduct Issues | 0.000 | 0.001 | -0.289 | 0.243 | 0.949 |
| Sex | 0.001 | 0.010 | 0.064 | 0.591 | 0.949 |
| **PPC positive** ($R^2$ = 0.068) | | | | | |
| CU traits | -0.002 | 0.002 | -0.789 | 0.365 | 0.573 |
| Tanner | 0.024 | 0.021 | 1.118 | 0.254 | 0.527 |
| Conduct Issues | -0.005 | 0.004 | -1.167 | 0.772 | 0.526 |
| Sex | 0.023 | 0.043 | 0.538 | 0.949 | 0.591 |
| **PPC negative** ($R^2$ = 0.031) | | | | | |
| CU traits | 0.000 | 0.000 | -0.031 | 0.976 | 0.975 |
| Tanner | 0.003 | 0.004 | 0.820 | 0.412 | 0.975 |
| Conduct Issues | 0.000 | 0.001 | 0.200 | 0.842 | 0.975 |
| Sex | 0.005 | 0.007 | 0.672 | 0.502 | 0.975 |

*= FDR p < 0.05

Supplementary Table 7. Within salience network centrality

| Network connections | β | S.E. | Z | p | FDR p |
|---|---|---|---|---|---|
| **ACC positive** ($R^2$ = 0.136) | | | | | |
| CU traits | 0.003 | 0.003 | 0.946 | 0.344 | 0.688 |
| Tanner | 0.015 | 0.025 | 0.602 | 0.547 | 0.729 |
| Conduct Issues | -0.012 | 0.005 | -2.499 | 0.012 | 0.049 |
| Sex | 0.008 | 0.050 | 0.152 | 0.880 | 0.879 |
| **ACC negative** ($R^2$ = 0.186) | | | | | |
| CU traits | 0.011 | 0.007 | 1.753 | 0.080 | 0.159 |
| Tanner | -0.033 | 0.056 | -0.593 | 0.553 | 0.737 |
| Conduct Issues | -0.027 | 0.010 | -2.630 | 0.009 | 0.034 |
| Sex | 0.020 | 0.112 | 0.178 | 0.859 | 0.858 |
| **Left insula positive (**$R^2$ = 0.202) | | | | | |
| CU traits | 0.010* | 0.003 | 3.026 | 0.002 | 0.009 |
| Tanner | 0.009 | 0.028 | 0.343 | 0.732 | 0.926 |
| Conduct Issues | 0.000 | 0.005 | 0.093 | 0.926 | 0.926 |
| Sex | 0.021 | 0.055 | 0.388 | 0.698 | 0.926 |
| **Left insula negative (**$R^2$ = 0.077) | | | | | |
| CU traits | -0.004 | 0.002 | -1.440 | 0.15 | 0.535 |
| Tanner | 0.018 | 0.021 | 0.843 | 0.399 | 0.535 |
| Conduct Issues | 0.004 | 0.004 | 1.077 | 0.282 | 0.535 |
| Sex | -0.008 | 0.042 | -0.185 | 0.853 | 0.853 |
| **Right insula positive** ($R^2$ = 0.124) | | | | | |
| CU traits | -0.006 | 0.003 | -1.811 | 0.07 | 0.170 |
| Tanner | -0.016 | 0.028 | -0.577 | 0.564 | 0.575 |
| Conduct Issues | 0.009 | 0.005 | 1.721 | 0.085 | 0.170 |
| Sex | 0.032 | 0.056 | 0.561 | 0.575 | 0.575 |
| **Right insula negative** ($R^2$ = 0.060) | | | | | |
| CU traits | -0.003 | 0.005 | -0.544 | 0.587 | 0.814 |
| Tanner | -0.011 | 0.045 | -0.234 | 0.815 | 0.814 |
| Conduct Issues | 0.013 | 0.009 | 1.539 | 0.124 | 0.495 |
| Sex | 0.044 | 0.091 | 0.482 | 0.63 | 0.814 |

*= FDR p < 0.05

Supplementary Table 8. Within frontoparietal network centrality

| Network connections | β | S.E. | Z | p | FDR p |
|---|---|---|---|---|---|
| **Left LPFC positive** ($R^2$ = 0.085) | | | | | |
| CU traits | 0.000 | 0.002 | -0.210 | 0.833 | 0.839 |
| Tanner | 0.009 | 0.017 | 0.513 | 0.608 | 0.839 |
| Conduct Issues | 0.006 | 0.003 | 1.792 | 0.073 | 0.293 |
| Sex | -0.007 | 0.035 | -0.202 | 0.84 | 0.839 |
| **Left LPFC negative** ($R^2$ = 0.033) | | | | | |
| CU traits | -0.002 | 0.002 | -0.992 | 0.321 | 0.789 |
| Tanner | 0.003 | 0.020 | 0.160 | 0.873 | 0.873 |
| Conduct Issues | 0.002 | 0.004 | 0.535 | 0.592 | 0.789 |
| Sex | -0.022 | 0.040 | -0.539 | 0.590 | 0.789 |
| **Left PCC positive** ($R^2$ = 0.105) | | | | | |
| CU traits | 0.003 | 0.002 | 1.605 | 0.108 | 0.433 |
| Tanner | 0.011 | 0.018 | 0.588 | 0.557 | 0.556 |
| Conduct Issues | -0.002 | 0.003 | -0.650 | 0.516 | 0.556 |
| Sex | 0.041 | 0.037 | 1.132 | 0.258 | 0.516 |
| **Left PCC negative** ($R^2$ = 0.140) | | | | | |
| CU traits | 0.008 | 0.004 | 2.147 | 0.032 | 0.127 |
| Tanner | 0.014 | 0.033 | 0.424 | 0.672 | 0.851 |
| Conduct Issues | -0.010 | 0.006 | -1.671 | 0.095 | 0.189 |
| Sex | 0.012 | 0.066 | 0.188 | 0.851 | 0.851 |
| **Right LPFC positive** ($R^2$ = 0.188) | | | | | |
| CU traits | -0.005* | 0.002 | -2.810 | 0.005 | 0.019 |
| Tanner | 0.003 | 0.017 | 0.190 | 0.849 | 0.849 |
| Conduct Issues | -0.001 | 0.003 | -0.471 | 0.637 | 0.849 |
| Sex | -0.027 | 0.033 | -0.800 | 0.424 | 0.847 |
| **Right LPFC negative** ($R^2$ = 0.210) | | | | | |
| CU traits | -0.010* | 0.003 | -3.151 | 0.002 | 0.006 |
| Tanner | -0.013 | 0.027 | -0.490 | 0.624 | 0.853 |
| Conduct Issues | 0.001 | 0.005 | 0.156 | 0.876 | 0.975 |
| Sex | 0.026 | 0.055 | 0.467 | 0.64 | 0.853 |
| **Right PCC positive** ($R^2$ = 0.059) | | | | | |
| CU traits | -0.002 | 0.004 | -0.662 | 0.508 | 0.929 |
| Tanner | 0.047 | 0.030 | 1.557 | 0.12 | 0.478 |
| Conduct Issues | 0.002 | 0.006 | 0.389 | 0.697 | 0.929 |
| Sex | -0.001 | 0.060 | -0.022 | 0.983 | 0.982 |
| **Right PCC negative** ($R^2$ = 0.080) | | | | | |
| CU traits | -0.003 | 0.005 | -0.544 | 0.725 | 0.725 |
| Tanner | -0.011 | 0.045 | -0.234 | 0.075 | 0.300 |
| Conduct Issues | 0.013 | 0.009 | 1.539 | 0.557 | 0.725 |
| Sex | 0.044 | 0.091 | 0.482 | 0.257 | 0.513 |

*= FDR $p < 0.05$

Supplementary Table 9. Positive between default mode and salience network centrality

| Network connections | β | S.E. | Z | p | FDR p |
|---|---|---|---|---|---|
| **mPFC-ACC positive** ($R^2$ = 0.024) | | | | | |
| CU traits | 0.000 | 0.000 | -0.95 | 0.342 | 0.684 |
| Tanner | -0.001 | 0.003 | -0.197 | 0.844 | 0.844 |
| Conduct Issues | 0.000 | 0.000 | 0.462 | 0.644 | 0.844 |
| Sex | 0.005 | 0.005 | 1.017 | 0.309 | 0.684 |
| **PCC-ACC positive** ($R^2$ = 0.106) | | | | | |
| CU traits | 0.003* | 0.001 | 2.912 | 0.004 | 0.014 |
| Tanner | 0.001 | 0.009 | 0.116 | 0.908 | 0.908 |
| Conduct Issues | -0.001 | 0.001 | -0.863 | 0.388 | 0.518 |
| Sex | 0.014 | 0.016 | 0.896 | 0.370 | 0.517 |
| **mPFC- left insula positive** ($R^2$ = 0.138) | | | | | |
| CU traits | 0.000 | 0.000 | 0.567 | 0.571 | 0.596 |
| Tanner | -0.008 | 0.002 | -3.485 | <0.001 | 0.002 |
| Conduct Issues | 0.000 | 0.000 | 0.884 | 0.377 | 0.596 |
| Sex | -0.002 | 0.004 | -0.529 | 0.597 | 0.597 |
| **PCC- left insula positive** ($R^2$ = 0.054) | | | | | |
| CU traits | -0.001 | 0.001 | -1.229 | 0.219 | 0.403 |
| Tanner | -0.006 | 0.009 | -0.695 | 0.487 | 0.487 |
| Conduct Issues | 0.002 | 0.002 | 1.594 | 0.111 | 0.403 |
| Sex | 0.017 | 0.017 | 1.03 | 0.303 | 0.404 |
| **mPFC- right insula positive** ($R^2$ = 0.023) | | | | | |
| CU traits | 0.000 | 0.001 | 0.318 | 0.751 | 0.751 |
| Tanner | 0.002 | 0.005 | 0.397 | 0.691 | 0.750 |
| Conduct Issues | 0.00 | 0.001 | 0.442 | 0.658 | 0.750 |
| Sex | -0.011 | 0.009 | -1.227 | 0.220 | 0.750 |
| **PCC- right insula positive** ($R^2$ = 0.037) | | | | | |
| CU traits | 0.00 | 0.00 | 0.418 | 0.676 | 0.676 |
| Tanner | -0.003 | 0.003 | -1.239 | 0.215 | 0.460 |
| Conduct Issues | 0.00 | 0.00 | -0.592 | 0.554 | 0.676 |
| Sex | 0.006 | 0.005 | 1.200 | 0.230 | 0.461 |

*= FDR p < 0.05

Supplementary Table 10. Negative between default mode and salience network centrality

| Network connections | β | S.E. | Z | p | FDR p |
|---|---|---|---|---|---|
| **mPFC-ACC negative** ($R^2$ = 0.064) | | | | | |
| CU traits | 0.000 | 0.002 | -0.210 | 0.833 | 0.855 |
| Tanner | 0.009 | 0.017 | 0.513 | 0.608 | 0.855 |
| Conduct Issues | 0.006 | 0.003 | 1.792 | 0.073 | 0.651 |
| Sex | -0.007 | 0.035 | -0.202 | 0.84 | 0.974 |
| **PCC-ACC negative** ($R^2$ = 0.159) | | | | | |
| CU traits | -0.002* | 0.002 | -0.992 | 0.321 | 0.024 |
| Tanner | 0.003 | 0.020 | 0.160 | 0.873 | 0.709 |
| Conduct Issues | 0.002 | 0.004 | 0.535 | 0.592 | 0.709 |
| Sex | -0.022 | 0.040 | -0.539 | 0.590 | 0.740 |
| **mPFC- left insula negative** ($R^2$ = 0.153) | | | | | |
| CU traits | 0.003 | 0.002 | 1.605 | 0.108 | 0.828 |
| Tanner | 0.011* | 0.018 | 0.588 | 0.557 | 0.024 |
| Conduct Issues | -0.002 | 0.003 | -0.650 | 0.516 | 0.811 |
| Sex | 0.041 | 0.037 | 1.132 | 0.258 | 0.265 |
| **PCC- left insula negative** ($R^2$ = 0.10) | | | | | |
| CU traits | 0.008 | 0.004 | 2.147 | 0.032 | 0.373 |
| Tanner | 0.014 | 0.033 | 0.424 | 0.672 | 0.888 |
| Conduct Issues | -0.010 | 0.006 | -1.671 | 0.095 | 0.312 |
| Sex | 0.012 | 0.066 | 0.188 | 0.851 | 0.695 |
| **mPFC- right insula negative** ($R^2$ = 0.024) | | | | | |
| CU traits | -0.005 | 0.002 | -2.810 | 0.005 | 0.844 |
| Tanner | 0.003 | 0.017 | 0.190 | 0.849 | 0.844 |
| Conduct Issues | -0.001 | 0.003 | -0.471 | 0.637 | 0.844 |
| Sex | -0.027 | 0.033 | -0.800 | 0.424 | 0.844 |
| **PCC- right insula negative** ($R^2$ = 0.062) | | | | | |
| CU traits | -0.010 | 0.003 | -3.151 | 0.002 | 0.683 |
| Tanner | -0.013 | 0.027 | -0.490 | 0.624 | 0.507 |
| Conduct Issues | 0.001 | 0.005 | 0.156 | 0.876 | 0.684 |
| Sex | 0.026 | 0.055 | 0.467 | 0.64 | 0.507 |

Note: some nodes were removed from analysis due to 0 connections.

*= FDR p < 0.05

Supplementary Table 11. Positive between default mode and frontoparietal network centrality

| Network connections | β | S.E. | Z | p | FDR p |
|---|---|---|---|---|---|
| **PCC-left LPFC positive** ($R^2$ = 0.187) | | | | | |
| CU traits | 0.002* | 0.001 | 4.144 | <0.001 | <0.001 |
| Tanner | -0.002 | 0.005 | -0.417 | 0.677 | 0.676 |
| Conduct Issues | -0.002 | 0.001 | -1.669 | 0.095 | 0.190 |
| Sex | 0.008 | 0.01 | 0.84 | 0.401 | 0.534 |
| **PCC- left PPC positive** ($R^2$ = 0.037) | | | | | |
| CU traits | 0.00 | 0.00 | -0.749 | 0.454 | 0.700 |
| Tanner | 0.001 | 0.004 | 0.162 | 0.872 | 0.871 |
| Conduct Issues | 0.001 | 0.001 | 1.555 | 0.120 | 0.479 |
| Sex | -0.005 | 0.007 | -0.635 | 0.525 | 0.701 |
| **PCC- right LPFC positive** ($R^2$ = 0.027) | | | | | |
| CU traits | 0.00 | 0.00 | -0.919 | 0.358 | 0.710 |
| Tanner | 0.001 | 0.002 | 0.371 | 0.711 | 0.711 |
| Conduct Issues | 0.00 | 0.00 | -0.478 | 0.633 | 0.710 |
| Sex | -0.004 | 0.003 | -1.057 | 0.290 | 0.710 |
| **PCC- right PPC positive** ($R^2$ = 0.093) | | | | | |
| CU traits | 0.001 | 0.001 | 2.17 | 0.030 | 0.114 |
| Tanner | 0.006 | 0.005 | 1.084 | 0.279 | 0.371 |
| Conduct Issues | -0.002 | 0.001 | -1.903 | 0.057 | 0.141 |
| Sex | -0.001 | 0.010 | -0.064 | 0.949 | 0.949 |

Note: some nodes were removed from analysis due to 0 connections.
*= FDR p < 0.05

Supplementary Table 12. Negative between default mode and frontoparietal network centrality

| Network connections | β | S.E. | Z | p | FDR p |
|---|---|---|---|---|---|
| **PCC-left LPFC positive** ($R^2$ = 0.166) | | | | | |
| CU traits | 0.003 | 0.001 | 2.456 | 0.014 | 0.056 |
| Tanner | 0.008 | 0.01 | 0.747 | 0.455 | 0.816 |
| Conduct Issues | 0.00 | 0.002 | 0.185 | 0.853 | 0.853 |
| Sex | 0.011 | 0.021 | 0.507 | 0.612 | 0.817 |
| **PCC- left PPC negative** ($R^2$ = 0.020) | | | | | |
| CU traits | 0.00 | 0.00 | -0.095 | 0.924 | 0.924 |
| Tanner | -0.001 | 0.003 | -0.177 | 0.86 | 0.924 |
| Conduct Issues | 0.00 | 0.001 | -0.331 | 0.741 | 0.924 |
| Sex | 0.005 | 0.006 | 0.875 | 0.381 | 0.924 |
| **mPFC- left PPC negative** ($R^2$ = 0.066) | | | | | |
| CU traits | -0.001 | 0.001 | -0.77 | 0.441 | 0.643 |
| Tanner | 0.003 | 0.009 | 0.306 | 0.76 | 0.759 |
| Conduct Issues | 0.003 | 0.002 | 1.431 | 0.152 | 0.610 |
| Sex | -0.013 | 0.019 | -0.702 | 0.483 | 0.643 |
| **PCC- left PPC negative** ($R^2$ = 0.041) | | | | | |
| CU traits | 0.00 | 0.00 | -0.939 | 0.348 | 0.696 |
| Tanner | 0.00 | 0.003 | -0.1 | 0.921 | 0.920 |
| Conduct Issues | 0.00 | 0.001 | 0.427 | 0.669 | 0.892 |
| Sex | 0.006 | 0.006 | 0.938 | 0.348 | 0.696 |
| **mPFC- right LPFC negative** ($R^2$ = 0.069) | | | | | |
| CU traits | 0.00 | 0.00 | 1.001 | 0.317 | 0.633 |
| Tanner | -0.007 | 0.004 | -1.588 | 0.112 | 0.448 |
| Conduct Issues | 0.00 | 0.001 | -0.399 | 0.69 | 0.919 |
| Sex | 0.00 | 0.009 | 0.021 | 0.984 | 0.984 |
| **PCC- right LPFC negative** ($R^2$ = 0.032) | | | | | |
| CU traits | 0.00 | 0.001 | 0.27 | 0.787 | 0.850 |
| Tanner | 0.001 | 0.005 | 0.188 | 0.851 | 0.850 |
| Conduct Issues | -0.001 | 0.001 | -0.703 | 0.482 | 0.850 |
| Sex | -0.01 | 0.011 | -0.953 | 0.34 | 0.850 |
| **PCC- right PPC negative** ($R^2$ = 0.225) | | | | | |
| CU traits | 0.004* | 0.001 | 2.817 | 0.005 | 0.019 |
| Tanner | 0.017 | 0.011 | 1.484 | 0.138 | 0.183 |
| Conduct Issues | -0.004 | 0.002 | -1.726 | 0.084 | 0.168 |
| Sex | -0.005 | 0.023 | -0.218 | 0.828 | 0.827 |

Note: some nodes were removed from analysis due to 0 connections.
*= FDR p < 0.05

Supplementary Table 13. Positive between salience and frontoparietal network centrality

| Network connections | β | S.E. | Z | p | FDR p |
|---|---|---|---|---|---|
| **Left Insula – left LPFC positive** ($R^2$ = 0.034) | | | | | |
| CU traits | 0.00 | 0.00 | -0.483 | 0.629 | 0.838 |
| Tanner | -0.002 | 0.002 | -1.233 | 0.217 | 0.452 |
| Conduct Issues | 0.00 | 0.00 | 0.087 | 0.93 | 0.930 |
| Sex | 0.004 | 0.003 | 1.211 | 0.226 | 0.452 |
| **Right Insula – left LPFC positive** ($R^2$ = 0.028) | | | | | |
| CU traits | 0.00 | 0.00 | 0.06 | 0.952 | 0.952 |
| Tanner | -0.001 | 0.003 | -0.349 | 0.727 | 0.952 |
| Conduct Issues | 0.001 | 0.001 | 1.367 | 0.171 | 0.685 |
| Sex | -0.004 | 0.006 | -0.668 | 0.504 | 0.952 |
| **ACC – Left PPC positive** ($R^2$ = 0.005) | | | | | |
| CU traits | 0.00 | 0.00 | -0.085 | 0.932 | 0.932 |
| Tanner | -0.001 | 0.005 | -0.208 | 0.835 | 0.932 |
| Conduct Issues | 0.00 | 0.001 | 0.365 | 0.715 | 0.932 |
| Sex | -0.004 | 0.009 | -0.492 | 0.622 | 0.932 |
| **Left Insula – left PPC positive (**$R^2$ = 0.037) | | | | | |
| CU traits | 0.00 | 0.00 | 0.418 | 0.676 | 0.676 |
| Tanner | -0.002 | 0.002 | -1.239 | 0.215 | 0.460 |
| Conduct Issues | 0.00 | 0.00 | -0.592 | 0.554 | 0.676 |
| Sex | 0.004 | 0.003 | 1.2 | 0.23 | 0.460 |
| **Right Insula – left PPC positive** ($R^2$ = 0.062) | | | | | |
| CU traits | 0.00 | 0.00 | -0.555 | 0.579 | 0.578 |
| Tanner | 0.005 | 0.004 | 1.489 | 0.136 | 0.298 |
| Conduct Issues | 0.001 | 0.001 | 1.086 | 0.278 | 0.370 |
| Sex | -0.009 | 0.007 | -1.442 | 0.149 | 0.298 |
| **ACC – right LPFC** ($R^2$ = 0.202) | | | | | |
| CU traits | 0.00 | 0.00 | -0.749 | 0.454 | 0.700 |
| Tanner | 0.001 | 0.004 | 0.162 | 0.872 | 0.871 |
| Conduct Issues | 0.001 | 0.001 | 1.555 | 0.12 | 0.479 |
| Sex | -0.005 | 0.007 | -0.635 | 0.525 | 0.700 |
| **Right Insula – right LPFC positive** ($R^2$ = 0.037) | | | | | |
| CU traits | 0.00 | 0.00 | 0.392 | 0.695 | 0.938 |
| Tanner | -0.002 | 0.003 | -0.866 | 0.386 | 0.932 |
| Conduct Issues | 0.00 | 0.00 | -0.184 | 0.854 | 0.932 |
| Sex | 0.00 | 0.005 | 0.084 | 0.933 | 0.932 |
| **ACC – right PPC positive** ($R^2$ = 0.011) | | | | | |
| CU traits | 0.00 | 0.00 | -0.918 | 0.358 | 0.716 |
| Tanner | 0.00 | 0.004 | 0.076 | 0.940 | 0.939 |
| Conduct Issues | 0.001 | 0.001 | 1.44 | 0.150 | 0.599 |
| Sex | -0.001 | 0.008 | -0.154 | 0.877 | 0.939 |
| **Left Insula – right PPC positive** ($R^2$ = 0.131) | | | | | |
| CU traits | -0.001 | 0.001 | -1.856 | 0.063 | 0.084 |
| Tanner | -0.014 | 0.007 | -1.907 | 0.057 | 0.084 |
| Conduct Issues | 0.003* | 0.001 | 2.729 | 0.006 | 0.025 |
| Sex | 0.007 | 0.013 | 0.546 | 0.585 | 0.585 |
| **Right Insula – right PPC positive** ($R^2$ = 0.062) | | | | | |
| CU traits | -0.001 | 0.001 | -1.159 | 0.246 | 0.371 |
| Tanner | -0.009 | 0.005 | -1.595 | 0.111 | 0.371 |
| Conduct Issues | 0.001 | 0.001 | 1.083 | 0.279 | 0.371 |
| Sex | -0.005 | 0.010 | -0.541 | 0.589 | 0.588 |

Note: some nodes were removed from analysis due to 0 connections.
*= FDR $p < 0.05$

Supplementary Table 14. Negative between salience and frontoparietal network centrality

| Network connections | β | S.E. | Z | p | FDR p |
|---|---|---|---|---|---|
| **ACC – left LPFC negative** ($R^2$ = 0.108) | | | | | |
| CU traits | 0.00 | 0.00 | 0.005 | 0.996 | 0.996 |
| Tanner | 0.006 | 0.004 | 1.408 | 0.159 | 0.220 |
| Conduct Issues | -0.001 | 0.001 | -1.386 | 0.166 | 0.221 |
| Sex | -0.014 | 0.008 | -1.666 | 0.096 | 0.220 |
| **Left Insula – left LPFC negative** ($R^2$ = 0.055) | | | | | |
| CU traits | 0.00 | 0.00 | -0.454 | 0.650 | 0.866 |
| Tanner | 0.00 | 0.00 | -1.146 | 0.252 | 0.503 |
| Conduct Issues | 0.00 | 0.00 | 0.023 | 0.982 | 0.981 |
| Sex | 0.001 | 0.001 | 1.17 | 0.242 | 0.503 |
| **Right Insula – left LPFC negative** ($R^2$ = 0.157) | | | | | |
| CU traits | 0.001 | 0.001 | 0.906 | 0.365 | 0.486 |
| Tanner* | -0.014 | 0.005 | -2.895 | 0.004 | 0.015 |
| Conduct Issues | 0.00 | 0.001 | -0.18 | 0.857 | 0.857 |
| Sex | 0.009 | 0.01 | 0.967 | 0.334 | 0.486 |
| **ACC – Left PPC negative** ($R^2$ = 0.016) | | | | | |
| CU traits | 0.00 | 0.001 | 0.213 | 0.831 | 0.990 |
| Tanner | 0.00 | 0.008 | -0.011 | 0.991 | 0.990 |
| Conduct Issues | 0.00 | 0.001 | 0.222 | 0.824 | 0.990 |
| Sex | -0.012 | 0.015 | -0.779 | 0.436 | 0.990 |
| **Left Insula – left PPC negative** ($R^2$ = 0.138) | | | | | |
| CU traits | 0.00 | 0.001 | -0.667 | 0.505 | 0.673 |
| Tanner | -0.013 | 0.005 | -2.396 | 0.017 | 0.066 |
| Conduct Issues | 0.00 | 0.001 | -0.187 | 0.852 | 0.851 |
| Sex | 0.012 | 0.011 | 1.075 | 0.283 | 0.565 |
| **Right Insula – left PPC negative** ($R^2$ = 0.036) | | | | | |
| CU traits | -0.001 | 0.001 | -1.107 | 0.269 | 0.881 |
| Tanner | -0.002 | 0.004 | -0.373 | 0.709 | 0.881 |
| Conduct Issues | 0.00 | 0.001 | 0.189 | 0.850 | 0.881 |
| Sex | -0.001 | 0.009 | -0.149 | 0.881 | 0.881 |
| **ACC – right LPFC negative** ($R^2$ = 0.110) | | | | | |
| CU traits | -0.001 | 0.001 | -1.316 | 0.188 | 0.252 |
| Tanner | 0.004 | 0.01 | 0.402 | 0.688 | 0.6875 |
| Conduct Issues | 0.003 | 0.002 | 1.467 | 0.142 | 0.252 |
| Sex | -0.025 | 0.019 | -1.311 | 0.190 | 0.252 |
| **Left Insula – right LPFC negative (**$R^2$ = 0.197) | | | | | |
| CU traits | 0.00 | 0.00 | 0.879 | 0.380 | 0.653 |
| Tanner | -0.009* | 0.003 | -3.318 | 0.001 | 0.003 |
| Conduct Issues | 0.00 | 0.001 | 0.69 | 0.490 | 0.650 |
| Sex | -0.002 | 0.006 | -0.419 | 0.675 | 0.675 |
| **Right Insula – right LPFC negative** ($R^2$ = 0.010) | | | | | |
| CU traits | 0.00 | 0.00 | 0.132 | 0.895 | 0.895 |
| Tanner | 0.00 | 0.00 | -0.629 | 0.529 | 0.529 |
| Conduct Issues | 0.00 | 0.00 | -0.137 | 0.891 | 0.891 |
| Sex | 0.00 | 0.001 | 0.219 | 0.827 | 0.826 |
| **ACC – right PPC negative** ($R^2$ = 0.070) | | | | | |
| CU traits | -0.002 | 0.001 | -1.37 | 0.171 | 0.560 |
| Tanner | 0.008 | 0.01 | 0.806 | 0.420 | 0.560 |
| Conduct Issues | 0.002 | 0.002 | 0.899 | 0.369 | 0.560 |
| Sex | -0.011 | 0.02 | -0.577 | 0.564 | 0.564 |
| **Left Insula – right PPC negative** ($R^2$ = 0.223) | | | | | |
| CU traits | -0.003 | 0.002 | -1.565 | 0.118 | 0.156 |
| Tanner | -0.028 | 0.015 | -1.849 | 0.065 | 0.129 |
| Conduct Issues | 0.009* | 0.003 | 3 | 0.003 | 0.010 |
| Sex | 0.011 | 0.031 | 0.369 | 0.712 | 0.712 |
| **Right Insula – right PPC negative** ($R^2$ = 0.052) | | | | | |
| CU traits | -0.001 | 0.001 | -0.777 | 0.437 | 0.583 |
| Tanner | -0.004 | 0.011 | -0.365 | 0.715 | 0.714 |
| Conduct Issues | 0.002 | 0.002 | 0.908 | 0.364 | 0.583 |
| Sex | -0.020 | 0.021 | -0.925 | 0.355 | 0.583 |

Note: some nodes were removed from analysis due to 0 connections.

*= FDR $p < 0.05$

Supplementary Table 15. Within network density – exploratory CU traits facets

| Network connections | B | S.E. | Z | P uncorrected |
|---|---|---|---|---|
| **DMN positive** ($R^2$ = 0.171) | | | | |
| Callousness | -0.002 | 0.002 | -0.897 | 0.370 |
| Uncaring | -0.001 | 0.001 | -1.020 | 0.308 |
| Unemotional | 0.002 | 0.002 | 0.782 | 0.434 |
| Tanner | 0.006 | 0.006 | 1.126 | 0.260 |
| Conduct Issues | -0.002* | 0.001 | -2.239 | 0.025 |
| Sex | -0.002 | 0.011 | -0.184 | 0.854 |
| **SAL positive** ($R^2$ = 0.108) | | | | |
| Callousness | -0.001 | 0.002 | -0.390 | 0.697 |
| Uncaring | 0.000 | 0.001 | -0.200 | 0.842 |
| Unemotional | 0.001 | 0.002 | 0.596 | 0.551 |
| Tanner | 0.006 | 0.006 | 0.936 | 0.349 |
| Conduct Issues | -0.002* | 0.001 | -2.066 | 0.039 |
| Sex | 0.003 | 0.012 | 0.214 | 0.831 |
| **FPN positive** ($R^2$ = 0.248) | | | | |
| Callousness | -0.002 | 0.001 | -1.478 | 0.139 |
| Uncaring | -0.001 | 0.001 | -1.819 | 0.069 |
| Unemotional | 0.001 | 0.001 | 0.648 | 0.517 |
| Tanner | 0.008 | 0.004 | 2.360 | 0.018 |
| Conduct Issues | -0.001 | 0.001 | -1.667 | 0.096 |
| Sex | -0.001 | 0.007 | -0.141 | 0.888 |
| **SAL negative** ($R^2$ = 0.113) | | | | |
| Callousness | -0.001 | 0.001 | -1.001 | 0.317 |
| Uncaring | 0.000 | 0.001 | -0.306 | 0.760 |
| Unemotional | 0.001 | 0.001 | 0.493 | 0.622 |
| Tanner | 0.001 | 0.003 | 0.218 | 0.828 |
| Conduct Issues | -0.001 | 0.001 | -1.687 | 0.092 |
| Sex | 0.003 | 0.007 | 0.406 | 0.685 |
| **FPN negative** ($R^2$ = 0.140) | | | | |
| Callousness | 0.000 | 0.001 | -0.038 | 0.969 |
| Uncaring | 0.000 | 0.001 | -0.707 | 0.479 |
| Unemotional | -0.001 | 0.001 | -0.872 | 0.383 |
| Tanner | -0.002 | 0.002 | -0.728 | 0.466 |
| Conduct Issues | -0.001 | 0.000 | -1.592 | 0.111 |
| Sex | 0.004 | 0.005 | 0.774 | 0.439 |

Note: DMN negative excluded because there were 0 negative connections
*= uncorrected $p < 0.05$

Supplementary Table 16. Between network density – exploratory CU traits facets

| Network connections | β | S.E. | Z | $P_{uncorrected}$ |
|---|---|---|---|---|
| **DMN-FPN positive** ($R^2$ = 0.195) | | | | |
| Callousness | 0.002 | 0.002 | 1.248 | 0.212 |
| Uncaring | 0.001 | 0.001 | 0.841 | 0.400 |
| Unemotional | 0.002 | 0.002 | 1.176 | 0.239 |
| Tanner | 0.004 | 0.005 | 0.824 | 0.410 |
| Conduct Issues | -0.001 | 0.001 | -1.180 | 0.238 |
| Sex | -0.006 | 0.010 | -0.657 | 0.511 |
| **DMN-FPN negative** ($R^2$ = 0.145) | | | | |
| Callousness | 0.001 | 0.002 | 0.457 | 0.647 |
| Uncaring | 0.002 | 0.001 | 1.095 | 0.274 |
| Unemotional | 0.003 | 0.002 | 1.117 | 0.264 |
| Tanner | 0.006 | 0.006 | 0.968 | 0.333 |
| Conduct Issues | 0.000 | 0.001 | -0.326 | 0.744 |
| Sex | 0.003 | 0.013 | 0.227 | 0.820 |
| **DMN-SAL positive** ($R^2$ = 0.194) | | | | |
| Callousness | 0.001 | 0.002 | 0.899 | 0.369 |
| Uncaring | 0.001 | 0.001 | 1.346 | 0.178 |
| Unemotional | -0.001 | 0.002 | -0.333 | 0.739 |
| Tanner | -0.006 | 0.005 | -1.388 | 0.165 |
| Conduct Issues | 0.002* | 0.001 | 2.045 | 0.041 |
| Sex | 0.007 | 0.009 | 0.758 | 0.448 |
| **DMN-SAL negative** ($R^2$ = 0.236) | | | | |
| Callousness | 0.001 | 0.002 | 0.294 | 0.769 |
| Uncaring | 0.002 | 0.001 | 1.619 | 0.105 |
| Unemotional | -0.002 | 0.002 | -1.166 | 0.244 |
| Tanner | -0.007 | 0.005 | -1.368 | 0.171 |
| Conduct Issues | 0.002* | 0.001 | 2.55 | 0.011 |
| Sex | 0.015 | 0.01 | 1.449 | 0.147 |
| **SAL-FPN positive** ($R^2$ = 0.216) | | | | |
| Callousness | 0.001 | 0.002 | 0.326 | 0.744 |
| Uncaring | -0.001 | 0.001 | -0.625 | 0.532 |
| Unemotional | -0.004 | 0.002 | -1.585 | 0.113 |
| Tanner | -0.006 | 0.006 | -0.915 | 0.360 |
| Conduct Issues | 0.003* | 0.001 | 2.549 | 0.011 |
| Sex | -0.014 | 0.013 | -1.155 | 0.248 |
| **SAL-FPN negative** ($R^2$ = 0.163) | | | | |
| Callousness | 0.00 | 0.002 | -0.045 | 0.964 |
| Uncaring | -0.001 | 0.001 | -0.856 | 0.392 |
| Unemotional | -0.002 | 0.002 | -0.896 | 0.370 |
| Tanner | -0.011 | 0.006 | -1.721 | 0.085 |
| Conduct Issues | 0.002 | 0.001 | 1.646 | 0.100 |
| Sex | -0.01 | 0.012 | -0.776 | 0.437 |

No ICU subscales are significant

Supplementary Table 17. Within default mode network centrality

| Network connections | β | S.E. | Z | P uncorrected |
|---|---|---|---|---|
| **mPFC positive** ($R^2$ = 0.085) | | | | |
| Callousness | -0.002 | 0.008 | -0.308 | 0.758 |
| Uncaring | 0.003 | 0.005 | 0.720 | 0.472 |
| Unemotional | 0.008 | 0.008 | 1.039 | 0.299 |
| Tanner | -0.019 | 0.021 | -0.883 | 0.377 |
| Conduct Issues | -0.001 | 0.004 | -0.253 | 0.800 |
| Sex | -0.037 | 0.043 | -0.869 | 0.385 |
| **mPFC negative** ($R^2$ = 0.042) | | | | |
| Callousness | 0.001 | 0.002 | 0.356 | 0.722 |
| Uncaring | 0.000 | 0.001 | 0.090 | 0.928 |
| Unemotional | 0.001 | 0.002 | 0.707 | 0.480 |
| Tanner | -0.005 | 0.005 | -1.074 | 0.283 |
| Conduct Issues | 0.000 | 0.001 | -0.297 | 0.767 |
| Sex | 0.002 | 0.010 | 0.163 | 0.870 |
| **PPC positive** ($R^2$ = 0.078) | | | | |
| Callousness | -0.006 | 0.008 | -0.816 | 0.414 |
| Uncaring | 0.000 | 0.005 | 0.047 | 0.962 |
| Unemotional | -0.001 | 0.008 | -0.135 | 0.893 |
| Tanner | 0.024 | 0.021 | 1.121 | 0.262 |
| Conduct Issues | -0.004 | 0.004 | -1.110 | 0.267 |
| Sex | 0.022 | 0.043 | 0.512 | 0.609 |
| **PPC negative** ($R^2$ = 0.031) | | | | |
| Callousness | 0.000 | 0.001 | -0.289 | 0.773 |
| Uncaring | 0.000 | 0.001 | 0.395 | 0.693 |
| Unemotional | 0.000 | 0.001 | -0.205 | 0.838 |
| Tanner | 0.003 | 0.004 | 0.784 | 0.433 |
| Conduct Issues | 0.000 | 0.001 | 0.229 | 0.819 |
| Sex | 0.004 | 0.008 | 0.596 | 0.551 |

No ICU subscales are significant

Supplementary Table 18. Within salience network centrality– exploratory CU traits facets

| Network connections | β | S.E. | Z | P uncorrected |
|---|---|---|---|---|
| **ACC positive** ($R^2$ = 0.136) | | | | |
| Callousness | 0.005 | 0.008 | -0.308 | 0.758 |
| Uncaring | 0.000 | 0.005 | 0.720 | 0.472 |
| Unemotional | 0.005 | 0.008 | 1.039 | 0.299 |
| Tanner | 0.016 | 0.021 | -0.883 | 0.377 |
| Conduct Issues | -0.012 | 0.004 | -0.253 | 0.800 |
| Sex | 0.012 | 0.043 | -0.869 | 0.385 |
| **ACC negative** ($R^2$ = 0.208) | | | | |
| Callousness | 0.015 | 0.008 | -0.816 | 0.414 |
| Uncaring | 0.001 | 0.005 | 0.047 | 0.962 |
| Unemotional | 0.030 | 0.008 | -0.135 | 0.893 |
| Tanner | -0.026 | 0.021 | 1.121 | 0.262 |
| Conduct Issues | -0.028 | 0.004 | -1.110 | 0.267 |
| Sex | 0.044 | 0.043 | 0.512 | 0.609 |
| **Left insula positive (**$R^2$ = 0.23) | | | | |
| Callousness | 0.007 | 0.002 | 0.356 | 0.722 |
| Uncaring | 0.013 | 0.001 | 0.090 | 0.928 |
| Unemotional | 0.005 | 0.002 | 0.707 | 0.480 |
| Tanner | 0.008 | 0.005 | -1.074 | 0.283 |
| Conduct Issues | 0.001 | 0.001 | -0.297 | 0.767 |
| Sex | 0.015 | 0.010 | 0.163 | 0.870 |
| **Left insula negative (**$R^2$ = 0.099) | | | | |
| Callousness | -0.011 | 0.001 | -0.289 | 0.773 |
| Uncaring | 0.002 | 0.001 | 0.395 | 0.693 |
| Unemotional | -0.005 | 0.001 | -0.205 | 0.838 |
| Tanner | 0.017 | 0.004 | 0.784 | 0.433 |
| Conduct Issues | 0.005 | 0.001 | 0.229 | 0.819 |
| Sex | -0.013 | 0.008 | 0.596 | 0.551 |
| **Right insula positive** ($R^2$ = 0.180) | | | | |
| Callousness | -0.010 | 0.009 | 0.604 | 0.546 |
| Uncaring | 0.003 | 0.006 | 0.002 | 0.998 |
| Unemotional | -0.019 | 0.010 | 0.565 | 0.572 |
| Tanner | -0.021 | 0.025 | 0.641 | 0.522 |
| Conduct Issues | 0.009 | 0.005 | -2.528 | 0.011 |
| Sex | 0.014 | 0.051 | 0.233 | 0.816 |
| **Right insula negative** ($R^2$ = 0.081) | | | | |
| Callousness | -0.011 | 0.010 | 0.715 | 0.475 |
| Uncaring | 0.007* | 0.006 | 2.129 | 0.033 |
| Unemotional | -0.014 | 0.011 | 0.516 | 0.606 |
| Tanner | -0.015 | 0.028 | 0.280 | 0.780 |
| Conduct Issues | 0.014 | 0.005 | 0.123 | 0.902 |
| Sex | 0.027 | 0.056 | 0.267 | 0.789 |

*= p < 0.05

Supplementary Table 19. Within frontoparietal network centrality– exploratory CU traits facets

| Network connections | β | S.E. | Z | P uncorrected |
|---|---|---|---|---|
| **Left LPFC positive** ($R^2$ = 0.093) | | | | |
| Callousness | -0.003 | 0.006 | -0.482 | 0.630 |
| Uncaring | -0.001 | 0.004 | -0.231 | 0.818 |
| Unemotional | 0.004 | 0.007 | 0.634 | 0.526 |
| Tanner | 0.010 | 0.017 | 0.603 | 0.546 |
| Conduct Issues | 0.006 | 0.003 | 1.833 | 0.067 |
| Sex | -0.003 | 0.035 | -0.075 | 0.941 |
| **Left LPFC negative** ($R^2$ = 0.085) | | | | |
| Callousness | 0.006 | 0.007 | 0.908 | 0.364 |
| Uncaring | -0.004 | 0.005 | -0.949 | 0.343 |
| Unemotional | -0.010 | 0.008 | -1.326 | 0.185 |
| Tanner | 0.001 | 0.020 | 0.037 | 0.970 |
| Conduct Issues | 0.002 | 0.004 | 0.423 | 0.672 |
| Sex | -0.027 | 0.040 | -0.673 | 0.501 |
| **Left PCC positive** ($R^2$ = 0.119) | | | | |
| Callousness | 0.001 | 0.007 | 0.083 | 0.934 |
| Uncaring | 0.004 | 0.004 | 0.839 | 0.402 |
| Unemotional | 0.007 | 0.007 | 1.046 | 0.296 |
| Tanner | 0.012 | 0.018 | 0.656 | 0.512 |
| Conduct Issues | -0.002 | 0.003 | -0.607 | 0.544 |
| Sex | 0.045 | 0.037 | 1.205 | 0.228 |
| **Left PCC negative** ($R^2$ = 0.139) | | | | |
| Callousness | 0.009 | 0.012 | 0.786 | 0.432 |
| Uncaring | 0.009 | 0.008 | 1.261 | 0.207 |
| Unemotional | 0.004 | 0.012 | 0.330 | 0.741 |
| Tanner | 0.012 | 0.033 | 0.376 | 0.707 |
| Conduct Issues | -0.010 | 0.006 | -1.672 | 0.094 |
| Sex | 0.008 | 0.067 | 0.118 | 0.906 |
| **Right LPFC positive** ($R^2$ = 0.239) | | | | |
| Callousness | -0.015* | 0.006 | -2.516 | 0.012 |
| Uncaring | 0.000 | 0.004 | -0.063 | 0.949 |
| Unemotional | -0.005 | 0.006 | -0.789 | 0.430 |
| Tanner | 0.003 | 0.016 | 0.188 | 0.851 |
| Conduct Issues | -0.001 | 0.003 | -0.326 | 0.744 |
| Sex | -0.030 | 0.033 | -0.909 | 0.364 |
| **Right LPFC negative** ($R^2$ = 0.209) | | | | |
| Callousness | -0.009 | 0.010 | -0.873 | 0.383 |
| Uncaring | -0.013* | 0.006 | -2.012 | 0.044 |
| Unemotional | -0.007 | 0.010 | -0.638 | 0.524 |
| Tanner | -0.012 | 0.028 | -0.439 | 0.661 |
| Conduct Issues | 0.001 | 0.005 | 0.138 | 0.890 |
| Sex | 0.031 | 0.056 | 0.546 | 0.585 |
| **Right PCC positive** ($R^2$ = 0.059) | | | | |
| Callousness | -0.005 | 0.011 | -0.510 | 0.610 |
| Uncaring | 0.000 | 0.007 | -0.058 | 0.954 |
| Unemotional | -0.002 | 0.011 | -0.204 | 0.838 |
| Tanner | 0.046 | 0.030 | 1.542 | 0.123 |
| Conduct Issues | 0.002 | 0.006 | 0.418 | 0.676 |
| Sex | -0.003 | 0.061 | -0.044 | 0.965 |
| **Right PCC negative** ($R^2$ = 0.093) | | | | |
| Callousness | 0.003 | 0.008 | 0.320 | 0.749 |
| Uncaring | 0.000 | 0.005 | 0.024 | 0.981 |
| Unemotional | -0.008 | 0.009 | -0.949 | 0.343 |
| Tanner | -0.043 | 0.022 | -1.891 | 0.059 |
| Conduct Issues | 0.002 | 0.004 | 0.548 | 0.583 |

| | | | | |
|---|---|---|---|---|
| Sex | 0.044 | 0.045 | 0.967 | 0.334 |

*= $p < 0.05$

Supplementary Table 20. Positive between default mode and salience network centrality – exploratory CU traits facets

| Network connections | β | S.E. | Z | $P_{uncorrected}$ |
|---|---|---|---|---|
| **mPFC-ACC positive** ($R^2$ = 0.049) | | | | |
| Callousness | 0.000 | 0.001 | -0.395 | 0.693 |
| Uncaring | -0.001 | 0.001 | -1.407 | 0.160 |
| Unemotional | 0.001 | 0.001 | 1.045 | 0.296 |
| Tanner | 0.000 | 0.003 | -0.041 | 0.967 |
| Conduct Issues | 0.000 | 0.000 | 0.426 | 0.670 |
| Sex | 0.006 | 0.005 | 1.203 | 0.229 |
| **PCC-ACC positive** ($R^2$ = 0.112) | | | | |
| Callousness | 0.005 | 0.003 | 1.554 | 0.120 |
| Uncaring | 0.002 | 0.002 | 0.984 | 0.325 |
| Unemotional | 0.002 | 0.003 | 0.570 | 0.569 |
| Tanner | 0.000 | 0.009 | 0.024 | 0.981 |
| Conduct Issues | -0.001 | 0.001 | -0.980 | 0.327 |
| Sex | 0.015 | 0.016 | 0.947 | 0.344 |
| **mPFC- left insula positive** ($R^2$ = 0.153) | | | | |
| Callousness | -0.001 | 0.001 | -0.864 | 0.388 |
| Uncaring | 0.000 | 0.001 | 0.492 | 0.623 |
| Unemotional | 0.001 | 0.001 | 1.105 | 0.269 |
| Tanner | -0.008 | 0.002 | -3.284 | 0.001 |
| Conduct Issues | 0.000 | 0.000 | 1.057 | 0.290 |
| Sex | -0.002 | 0.004 | -0.550 | 0.583 |
| **PCC- left insula positive** ($R^2$ = 0.064) | | | | |
| Callousness | -0.003 | 0.003 | -0.998 | 0.318 |
| Uncaring | 0.001 | 0.002 | 0.248 | 0.804 |
| Unemotional | -0.002 | 0.003 | -0.680 | 0.496 |
| Tanner | -0.006 | 0.009 | -0.672 | 0.502 |
| Conduct Issues | 0.003 | 0.002 | 1.719 | 0.086 |
| Sex | 0.015 | 0.017 | 0.877 | 0.381 |
| **mPFC- right insula positive** ($R^2$ = 0.037) | | | | |
| Callousness | 0.001 | 0.002 | 0.419 | 0.675 |
| Uncaring | -0.001 | 0.001 | -0.772 | 0.440 |
| Unemotional | 0.002 | 0.002 | 0.883 | 0.377 |
| Tanner | 0.002 | 0.005 | 0.455 | 0.649 |
| Conduct Issues | 0.000 | 0.001 | 0.335 | 0.737 |
| Sex | -0.010 | 0.009 | -1.050 | 0.294 |
| **PCC- right insula positive** ($R^2$ = 0.061) | | | | |
| Callousness | 0.001 | 0.001 | 1.170 | 0.242 |
| Uncaring | 0.000 | 0.001 | 0.253 | 0.800 |
| Unemotional | -0.001 | 0.001 | -1.169 | 0.242 |
| Tanner | -0.004 | 0.003 | -1.469 | 0.142 |
| Conduct Issues | 0.000 | 0.000 | -0.757 | 0.449 |
| Sex | 0.006 | 0.005 | 1.168 | 0.243 |

No ICU subscales are significant

Supplementary Table 21. Negative between default mode and salience network centrality – exploratory CU traits facets

| Network connections | β | S.E. | Z | $P_{uncorrected}$ |
|---|---|---|---|---|
| **mPFC-ACC negative** ($R^2$ = 0.099) | | | | |
| Callousness | -0.001 | 0.003 | -0.345 | 0.730 |
| Uncaring | 0.000 | 0.002 | 0.124 | 0.901 |
| Unemotional | 0.005 | 0.003 | 1.419 | 0.156 |
| Tanner | -0.003 | 0.009 | -0.314 | 0.753 |
| Conduct Issues | 0.002 | 0.002 | 1.468 | 0.142 |
| Sex | 0.005 | 0.018 | 0.249 | 0.804 |
| **PCC-ACC negative** ($R^2$ = 0.161) | | | | |
| Callousness | 0.008 | 0.005 | 1.565 | 0.118 |
| Uncaring | 0.004 | 0.003 | 1.213 | 0.225 |
| Unemotional | 0.001 | 0.006 | 0.252 | 0.801 |
| Tanner | -0.011 | 0.015 | -0.697 | 0.486 |
| Conduct Issues | -0.002 | 0.003 | -0.831 | 0.406 |
| Sex | 0.008 | 0.031 | 0.245 | 0.807 |
| **mPFC- left insula negative** ($R^2$ = 0.152) | | | | |
| Callousness | 0.000 | 0.001 | 0.222 | 0.824 |
| Uncaring | 0.000 | 0.001 | -0.361 | 0.718 |
| Unemotional | 0.000 | 0.002 | -0.106 | 0.915 |
| Tanner | -0.011 | 0.004 | -2.726 | 0.006 |
| Conduct Issues | 0.000 | 0.001 | 0.481 | 0.631 |
| Sex | 0.012 | 0.008 | 1.490 | 0.136 |
| **PCC- left insula negative** ($R^2$ = 0.145) | | | | |
| Callousness | -0.006 | 0.006 | -0.933 | 0.351 |
| Uncaring | 0.002 | 0.004 | 0.595 | 0.552 |
| Unemotional | -0.010 | 0.007 | -1.524 | 0.127 |
| Tanner | -0.005 | 0.018 | -0.304 | 0.761 |
| Conduct Issues | 0.006 | 0.003 | 1.859 | 0.063 |
| Sex | 0.013 | 0.036 | 0.360 | 0.719 |
| **mPFC- right insula negative** ($R^2$ = 0.040) | | | | |
| Callousness | 0.000 | 0.003 | 0.073 | 0.942 |
| Uncaring | 0.001 | 0.002 | 0.407 | 0.684 |
| Unemotional | -0.003 | 0.004 | -0.924 | 0.355 |
| Tanner | 0.001 | 0.009 | 0.102 | 0.919 |
| Conduct Issues | 0.001 | 0.002 | 0.675 | 0.500 |
| Sex | 0.008 | 0.019 | 0.428 | 0.669 |
| **PCC- right insula negative** ($R^2$ = 0.092) | | | | |
| Callousness | 0.000 | 0.000 | 0.970 | 0.332 |
| Uncaring | 0.000 | 0.000 | 0.403 | 0.687 |
| Unemotional | 0.000 | 0.000 | -1.205 | 0.228 |
| Tanner | -0.001 | 0.000 | -1.456 | 0.145 |
| Conduct Issues | 0.000 | 0.000 | -0.629 | 0.529 |
| Sex | 0.001 | 0.001 | 0.908 | 0.364 |

Note: some nodes were removed from analysis due to 0 connections.
No ICU subscales are significant

Supplementary Table 22. Positive between default mode and frontoparietal network centrality – exploratory CU traits facets

| Network connections | β | S.E. | Z | P uncorrected |
|---|---|---|---|---|
| **PCC-left LPFC positive** ($R^2$ = 0.206) | | | | |
| Callousness | 0.005* | 0.002 | 2.604 | 0.009 |
| Uncaring | 0.001 | 0.001 | 1.044 | 0.296 |
| Unemotional | 0.002 | 0.002 | 0.845 | 0.398 |
| Tanner | -0.003 | 0.005 | -0.575 | 0.565 |
| Conduct Issues | -0.002 | 0.001 | -1.926 | 0.054 |
| Sex | 0.010 | 0.010 | 0.988 | 0.323 |
| **PCC- left PPC positive** ($R^2$ = 0.042) | | | | |
| Callousness | 0.000 | 0.001 | -0.216 | 0.829 |
| Uncaring | -0.001 | 0.001 | -0.824 | 0.410 |
| Unemotional | 0.000 | 0.001 | 0.360 | 0.719 |
| Tanner | 0.001 | 0.004 | 0.224 | 0.823 |
| Conduct Issues | 0.001 | 0.001 | 1.504 | 0.133 |
| Sex | -0.004 | 0.007 | -0.538 | 0.590 |
| **PCC- right LPFC positive** ($R^2$ = 0.068) | | | | |
| Callousness | -0.001 | 0.001 | -1.276 | 0.202 |
| Uncaring | 0.000 | 0.000 | -0.947 | 0.344 |
| Unemotional | 0.001 | 0.001 | 1.528 | 0.126 |
| Tanner | 0.001 | 0.002 | 0.656 | 0.512 |
| Conduct Issues | 0.000 | 0.000 | -0.344 | 0.731 |
| Sex | -0.003 | 0.003 | -0.948 | 0.343 |
| **PCC- right PPC positive** ($R^2$ = 0.094) | | | | |
| Callousness | 0.001 | 0.002 | 0.451 | 0.652 |
| Uncaring | 0.001 | 0.001 | 0.918 | 0.358 |
| Unemotional | 0.002 | 0.002 | 1.015 | 0.310 |
| Tanner | 0.006 | 0.006 | 1.128 | 0.259 |
| Conduct Issues | -0.002 | 0.001 | -1.842 | 0.066 |
| Sex | 0.000 | 0.010 | -0.041 | 0.967 |

Note: some nodes were removed from analysis due to 0 connections.

*= $p < 0.05$

Supplementary Table 23. Negative between default mode and frontoparietal network centrality – exploratory CU traits facets

| Network connections | β | S.E. | Z | $P_{uncorrected}$ |
|---|---|---|---|---|
| **PCC-left LPFC positive** ($R^2$ = 0.157) | | | | |
| Callousness | 0.001 | 0.004 | 0.212 | 0.832 |
| Uncaring | 0.005 | 0.002 | 2.113 | 0.035 |
| Unemotional | 0.002 | 0.004 | 0.390 | 0.697 |
| Tanner | 0.007 | 0.010 | 0.689 | 0.491 |
| Conduct Issues | 0.000 | 0.002 | 0.249 | 0.803 |
| Sex | 0.008 | 0.021 | 0.373 | 0.709 |
| **PCC- left PPC negative** ($R^2$ = 0.037) | | | | |
| Callousness | 0.000 | 0.001 | -0.001 | 0.999 |
| Uncaring | 0.000 | 0.001 | 0.536 | 0.592 |
| Unemotional | -0.001 | 0.001 | -0.860 | 0.390 |
| Tanner | -0.001 | 0.003 | -0.292 | 0.770 |
| Conduct Issues | 0.000 | 0.001 | -0.331 | 0.740 |
| Sex | 0.004 | 0.006 | 0.683 | 0.495 |
| **mPFC- left PPC negative** ($R^2$ = 0.077) | | | | |
| Callousness | -0.001 | 0.003 | -0.223 | 0.824 |
| Uncaring | -0.002 | 0.002 | -0.905 | 0.365 |
| Unemotional | 0.001 | 0.004 | 0.410 | 0.682 |
| Tanner | 0.004 | 0.009 | 0.396 | 0.692 |
| Conduct Issues | 0.003 | 0.002 | 1.427 | 0.153 |
| Sex | -0.010 | 0.019 | -0.546 | 0.585 |
| **PCC- left PPC negative** ($R^2$ = 0.078) | | | | |
| Callousness | 0.000 | 0.001 | -0.451 | 0.652 |
| Uncaring | -0.001 | 0.001 | -1.263 | 0.207 |
| Unemotional | 0.001 | 0.001 | 0.930 | 0.353 |
| Tanner | 0.000 | 0.003 | 0.068 | 0.946 |
| Conduct Issues | 0.000 | 0.001 | 0.442 | 0.658 |
| Sex | 0.007 | 0.006 | 1.202 | 0.229 |
| **mPFC- right LPFC negative** ($R^2$ = 0.116) | | | | |
| Callousness | 0.000 | 0.001 | -0.137 | 0.891 |
| Uncaring | 0.000 | 0.001 | -0.070 | 0.944 |
| Unemotional | 0.003 | 0.002 | 1.724 | 0.085 |
| Tanner | -0.006 | 0.004 | -1.422 | 0.155 |
| Conduct Issues | 0.000 | 0.001 | -0.366 | 0.715 |
| Sex | 0.003 | 0.008 | 0.301 | 0.763 |
| **PCC- right LPFC negative** ($R^2$ = 0.041) | | | | |
| Callousness | 0.001 | 0.002 | 0.289 | 0.773 |
| Uncaring | -0.001 | 0.001 | -0.442 | 0.658 |
| Unemotional | 0.001 | 0.002 | 0.610 | 0.542 |
| Tanner | 0.001 | 0.005 | 0.264 | 0.792 |
| Conduct Issues | -0.001 | 0.001 | -0.727 | 0.467 |
| Sex | -0.009 | 0.011 | -0.808 | 0.419 |
| **PCC- right PPC negative** ($R^2$ = 0.405) | | | | |
| Callousness | 0.003 | 0.004 | 0.702 | 0.483 |
| Uncaring | 0.003 | 0.003 | 1.150 | 0.250 |
| Unemotional | 0.007 | 0.004 | 1.558 | 0.119 |
| Tanner | 0.018 | 0.011 | 1.573 | 0.116 |
| Conduct Issues | -0.004 | 0.002 | -1.708 | 0.088 |
| Sex | -0.002 | 0.023 | -0.077 | 0.938 |

Note: some nodes were removed from analysis due to 0 connections.
No ICU subscales are significant

Supplementary Table 24. Positive between salience and frontoparietal network centrality – exploratory CU traits facets

| Network connections | β | S.E. | Z | $P_{uncorrected}$ |
|---|---|---|---|---|
| **Left Insula – left LPFC positive** ($R^2$ = 0.066) | | | | |
| Callousness | 0.000 | 0.001 | 0.713 | 0.476 |
| Uncaring | -0.001 | 0.000 | -1.709 | 0.088 |
| Unemotional | 0.000 | 0.001 | 0.711 | 0.477 |
| Tanner | -0.002 | 0.002 | -1.216 | 0.224 |
| Conduct Issues | 0.000 | 0.000 | -0.133 | 0.894 |
| Sex | 0.005 | 0.003 | 1.483 | 0.138 |
| **Right Insula – left LPFC positive** ($R^2$ = 0.031) | | | | |
| Callousness | 0.000 | 0.001 | -0.257 | 0.797 |
| Uncaring | 0.000 | 0.001 | 0.504 | 0.615 |
| Unemotional | 0.000 | 0.001 | -0.260 | 0.795 |
| Tanner | -0.001 | 0.003 | -0.353 | 0.724 |
| Conduct Issues | 0.001 | 0.001 | 1.415 | 0.157 |
| Sex | -0.004 | 0.006 | -0.747 | 0.455 |
| **ACC – Left PPC positive** ($R^2$ = 0.055) | | | | |
| Callousness | 0.003 | 0.002 | 1.860 | 0.063 |
| Uncaring | -0.001 | 0.001 | -0.677 | 0.498 |
| Unemotional | -0.002 | 0.002 | -1.438 | 0.150 |
| Tanner | -0.002 | 0.005 | -0.523 | 0.601 |
| Conduct Issues | 0.000 | 0.001 | 0.025 | 0.980 |
| Sex | -0.004 | 0.009 | -0.415 | 0.678 |
| **Left Insula – left PPC positive (**$R^2$ = 0.061) | | | | |
| Callousness | 0.001 | 0.001 | 1.170 | 0.242 |
| Uncaring | 0.000 | 0.000 | 0.253 | 0.800 |
| Unemotional | -0.001 | 0.001 | -1.169 | 0.242 |
| Tanner | -0.003 | 0.002 | -1.469 | 0.142 |
| Conduct Issues | 0.000 | 0.000 | -0.757 | 0.449 |
| Sex | 0.004 | 0.003 | 1.168 | 0.243 |
| **Right Insula – left PPC positive** ($R^2$ = 0.083) | | | | |
| Callousness | 0.001 | 0.001 | 0.428 | 0.668 |
| Uncaring | 0.000 | 0.001 | 0.207 | 0.836 |
| Unemotional | -0.002 | 0.001 | -1.465 | 0.143 |
| Tanner | 0.005 | 0.004 | 1.295 | 0.195 |
| Conduct Issues | 0.001 | 0.001 | 1.004 | 0.315 |
| Sex | -0.010 | 0.007 | -1.541 | 0.123 |
| **ACC – right LPFC** ($R^2$ = 0.042) | | | | |
| Callousness | 0.000 | 0.001 | -0.216 | 0.829 |
| Uncaring | -0.001 | 0.001 | -0.824 | 0.410 |
| Unemotional | 0.000 | 0.001 | 0.360 | 0.719 |
| Tanner | 0.001 | 0.004 | 0.224 | 0.823 |
| Conduct Issues | 0.001 | 0.001 | 1.504 | 0.133 |
| Sex | -0.004 | 0.007 | -0.538 | 0.590 |
| **Right Insula – right LPFC positive** ($R^2$ = 0.016) | | | | |
| Callousness | 0.000 | 0.001 | -0.266 | 0.790 |
| Uncaring | 0.000 | 0.001 | 0.064 | 0.949 |
| Unemotional | 0.001 | 0.001 | 0.721 | 0.471 |
| Tanner | -0.002 | 0.003 | -0.760 | 0.448 |
| Conduct Issues | 0.000 | 0.000 | -0.124 | 0.901 |
| Sex | 0.001 | 0.005 | 0.114 | 0.909 |
| **ACC – right PPC positive** ($R^2$ = 0.037) | | | | |
| Callousness | 0.000 | 0.002 | -0.234 | 0.815 |
| Uncaring | -0.001 | 0.001 | -0.922 | 0.357 |
| Unemotional | 0.000 | 0.002 | 0.294 | 0.769 |
| Tanner | 0.001 | 0.004 | 0.134 | 0.893 |
| Conduct Issues | 0.001 | 0.001 | 1.384 | 0.166 |

| | | | | |
|---|---|---|---|---|
| Sex | 0.000 | 0.008 | -0.059 | 0.953 |
| **Left Insula – right PPC positive** ($R^2$ = 0.136) | | | | |
| Callousness | -0.003 | 0.003 | -1.244 | 0.213 |
| Uncaring | -0.001 | 0.002 | -0.396 | 0.692 |
| Unemotional | -0.001 | 0.003 | -0.357 | 0.721 |
| Tanner | -0.013 | 0.007 | -1.810 | 0.070 |
| Conduct Issues | 0.004 | 0.001 | 2.825 | 0.005 |
| Sex | 0.006 | 0.013 | 0.463 | 0.643 |
| **Right Insula – right PPC positive** ($R^2$ = 0.066) | | | | |
| Callousness | -0.001 | 0.002 | -0.368 | 0.713 |
| Uncaring | -0.001 | 0.001 | -0.925 | 0.355 |
| Unemotional | 0.000 | 0.002 | 0.155 | 0.877 |
| Tanner | -0.008 | 0.005 | -1.521 | 0.128 |
| Conduct Issues | 0.001 | 0.001 | 1.048 | 0.295 |
| Sex | -0.005 | 0.010 | -0.461 | 0.645 |

Note: some nodes were removed from analysis due to 0 connections.
No ICU subscales are significant

Supplementary Table 25. Negative between salience and frontoparietal network centrality – exploratory CU traits facets

| Network connections | β | S.E. | Z | $P_{uncorrected}$ |
|---|---|---|---|---|
| **ACC – left LPFC negative** ($R^2$ = 0.144) | | | | |
| Callousness | 0.000 | 0.001 | -0.129 | 0.898 |
| Uncaring | -0.001 | 0.001 | -0.887 | 0.375 |
| Unemotional | 0.002 | 0.002 | 1.370 | 0.171 |
| Tanner | 0.007 | 0.004 | 1.620 | 0.105 |
| Conduct Issues | -0.001 | 0.001 | -1.408 | 0.159 |
| Sex | -0.011 | 0.008 | -1.378 | 0.168 |
| **Left Insula – left LPFC negative** ($R^2$ = 0.087) | | | | |
| Callousness | 0.000 | 0.000 | 0.528 | 0.598 |
| Uncaring | 0.000 | 0.000 | -1.444 | 0.149 |
| Unemotional | 0.000 | 0.000 | 0.612 | 0.541 |
| Tanner | 0.000 | 0.000 | -1.047 | 0.295 |
| Conduct Issues | 0.000 | 0.000 | -0.049 | 0.961 |
| Sex | 0.001 | 0.001 | 1.395 | 0.163 |
| **Right Insula – left LPFC negative** ($R^2$ = 0.167) | | | | |
| Callousness | 0.000 | 0.002 | 0.096 | 0.923 |
| Uncaring | 0.000 | 0.001 | 0.238 | 0.812 |
| Unemotional | 0.002 | 0.002 | 0.844 | 0.399 |
| Tanner* | -0.014 | 0.005 | -2.806 | 0.005 |
| Conduct Issues | 0.000 | 0.001 | -0.162 | 0.871 |
| Sex | 0.011 | 0.010 | 1.064 | 0.287 |
| **ACC – Left PPC negative** ($R^2$ = 0.016) | | | | |
| Callousness | 0.000 | 0.003 | 0.140 | 0.889 |
| Uncaring | 0.000 | 0.002 | 0.026 | 0.979 |
| Unemotional | 0.000 | 0.003 | 0.086 | 0.931 |
| Tanner | 0.000 | 0.008 | -0.007 | 0.994 |
| Conduct Issues | 0.000 | 0.001 | 0.214 | 0.831 |
| Sex | -0.012 | 0.016 | -0.754 | 0.451 |
| **Left Insula – left PPC negative** ($R^2$ = 0.165) | | | | |
| Callousness | 0.002 | 0.002 | 1.042 | 0.297 |
| Uncaring | -0.001 | 0.001 | -0.848 | 0.396 |
| Unemotional | -0.002 | 0.002 | -1.164 | 0.244 |
| Tanner | -0.014 | 0.005 | -2.542 | 0.011 |
| Conduct Issues | 0.000 | 0.001 | -0.314 | 0.753 |

| | | | | |
|---|---|---|---|---|
| Sex | 0.010 | 0.011 | 0.959 | 0.338 |
| **Right Insula – left PPC negative** ($R^2$ = 0.112) | | | | |
| Callousness | -0.002 | 0.001 | -1.407 | 0.160 |
| Uncaring | -0.001 | 0.001 | -1.090 | 0.276 |
| Unemotional | 0.003 | 0.002 | 1.645 | 0.100 |
| Tanner | -0.001 | 0.004 | -0.122 | 0.903 |
| Conduct Issues | 0.000 | 0.001 | 0.292 | 0.771 |
| Sex | 0.002 | 0.008 | 0.217 | 0.828 |
| **ACC – right LPFC negative** ($R^2$ = 0.116) | | | | |
| Callousness | 0.000 | 0.003 | -0.129 | 0.897 |
| Uncaring | -0.002 | 0.002 | -0.825 | 0.409 |
| Unemotional | -0.002 | 0.004 | -0.587 | 0.557 |
| Tanner | 0.004 | 0.010 | 0.378 | 0.705 |
| Conduct Issues | 0.003 | 0.002 | 1.433 | 0.152 |
| Sex | -0.026 | 0.020 | -1.306 | 0.191 |
| **Left Insula – right LPFC negative (**$R^2$ = 0.230) | | | | |
| Callousness | -0.001 | 0.001 | -0.926 | 0.354 |
| Uncaring | 0.001 | 0.001 | 1.003 | 0.316 |
| Unemotional | 0.001 | 0.001 | 1.109 | 0.267 |
| Tanner | -0.009 | 0.003 | -3.253 | 0.001 |
| Conduct Issues | 0.000 | 0.001 | 0.820 | 0.412 |
| Sex | -0.002 | 0.006 | -0.325 | 0.745 |
| **Right Insula – right LPFC negative** ($R^2$ = 0.015) | | | | |
| Callousness | 0.000 | 0.000 | -0.298 | 0.766 |
| Uncaring | 0.000 | 0.000 | 0.119 | 0.905 |
| Unemotional | 0.000 | 0.000 | 0.410 | 0.681 |
| Tanner | 0.000 | 0.000 | -0.578 | 0.563 |
| Conduct Issues | 0.000 | 0.000 | -0.104 | 0.918 |
| Sex | 0.000 | 0.001 | 0.272 | 0.786 |
| **ACC – right PPC negative** ($R^2$ = 0.098) | | | | |
| Callousness | 0.002 | 0.004 | 0.471 | 0.637 |
| Uncaring | -0.003 | 0.002 | -1.266 | 0.206 |
| Unemotional | -0.003 | 0.004 | -0.861 | 0.389 |
| Tanner | 0.008 | 0.010 | 0.760 | 0.447 |
| Conduct Issues | 0.002 | 0.002 | 0.812 | 0.417 |
| Sex | -0.012 | 0.020 | -0.599 | 0.549 |
| **Left Insula – right PPC negative** ($R^2$ = 0.253) | | | | |
| Callousness | -0.002 | 0.005 | -0.454 | 0.650 |
| Uncaring | 0.000 | 0.003 | -0.042 | 0.967 |
| Unemotional | -0.009 | 0.006 | -1.616 | 0.106 |
| Tanner | -0.031 | 0.015 | -2.022 | 0.043 |
| Conduct Issues | 0.009 | 0.003 | 3.038 | 0.002 |
| Sex | 0.004 | 0.031 | 0.120 | 0.904 |
| **Right Insula – right PPC negative** ($R^2$ = 0.055) | | | | |
| Callousness | -0.003 | 0.004 | -0.667 | 0.505 |
| Uncaring | 0.000 | 0.002 | 0.016 | 0.987 |
| Unemotional | -0.001 | 0.004 | -0.265 | 0.791 |
| Tanner | -0.004 | 0.011 | -0.373 | 0.709 |
| Conduct Issues | 0.002 | 0.002 | 0.948 | 0.343 |
| Sex | -0.021 | 0.022 | -0.948 | 0.343 |

Note: some nodes were removed from analysis due to 0 connections.
No ICU subscales are significant
